# Multi-lineage transcriptional and cell communication signatures define evolving personalized mechanisms that initiate and perpetuate rheumatoid arthritis

**DOI:** 10.1101/2025.02.08.619913

**Authors:** Cong Liu, Z. Audrey Wang, E. Barton Prideaux, David L Boyle, Katherine Nguyen, Vlad Tsaltskan, Amy Westermann, Andrea Ochoa, Kevin D. Deane, Marie L. Feser, M. Kristen Demoruelle, Kristine A. Kuhn, V. Michael Holers, Fan Zhang, Ashley G. Asamoah, Lauren Okada, Mark A. Gillespie, Palak Genge, Morgan Weiss, Veronica Hernandez, Julian Reading, Lynne Becker, Jane H. Buckner, Cate Speake, Jieyuan Liu, Zhen Wang, Zhiting Hu, Thomas F. Bumol, Peter Skene, Gary S. Firestein, Wei Wang

## Abstract

Rheumatoid arthritis (RA) is a systemic autoimmune disease arising from loss of tolerance and autoantibody development in at-risk individuals. Targeted therapies yield variable responses and current preventive strategies delay but do not stop disease onset. We propose that a set of transcription factors (TFs) and their downstream pathways regulate inflammatory cell communication networks in at-risk populations and RA. These networks enable multiple pathogenic cell types and mediators and could account for variable responses to targeted agents. To test this hypothesis, we identified anti-citrullinated protein antibody (ACPA)-positive at-risk individuals, patients with early and established RA and healthy controls. Single cell chromatin accessibility and transcriptomic profiles from blood mononuclear r cells were integrated and identified share pathogenic mechanisms, especially SUMOylation, RUNX2, YAP1, NOTCH3, and β-Catenin Pathways. Surprisingly, this signature was found in multiple cell types. Individualized gene expression patterns were then confirmed in RA synovium. Cell communication analysis revealed that multiple lineages can deliver a core set of pro-inflammatory mediators to receiver cells. Longitudinal analysis showed that the signature cell types evolve in individual at-risk participants. Cell-type-specific signature pathways could contribute to the differing clinical responses to targeted therapies. This study describes how a common clinical phenotype could arise from multiple pathogenic mechanisms.

## Introduction

Rheumatoid arthritis (RA) is a systemic immune-mediated disease marked by synovial inflammation and joint destruction^1^. While recent advances led to targeted therapies that block specific cytokines or immune cell types, clinical responses are highly variable. A lack of response to one targeted agent does not preclude a response to another with a different mechanism of action.

Current models suggest that seropositive RA begins with mucosal inflammation and loss of self-tolerance in individuals that carry genetic risk alleles and are exposed to risk-elevating environmental factors^2^. During a prolonged asymptomatic phase, anti-citrullinated protein antibody (ACPA) levels increase. The presence of these autoantibodies is strongly associated with the future development of RA in up to 60% of at-risk individuals^3^. While several clinical trials have attempted to prevent onset of synovitis, including treatment with atorvastatin, rituximab, methotrexate, hydroxychloroquine, and abatacept^4–8^, they have not been shown to prevent RA.

These observations led us to propose that diverse mechanisms in individuals at risk for developing RA and with RA ultimately converge to produce a common clinical phenotype^1^. However, these divergent pathogenic pathways are poorly understood, limiting our ability to predict the benefits of targeted therapeutics for individual patients. A challenging question is: how do heterogeneous mechanisms in at-risk individuals converge to create a common phenotype? We hypothesized that a unifying set of transcription factors (TFs) and their downstream genes and pathways regulate a pro-inflammatory cell communication network and enabling multiple diverse cell types to serve as pathogenic drivers in different individuals^1^. In this model, pathogenesis is defined by a shared transcriptional program, even if the specific cell types and inflammatory mediators initiating the synovitis vary from patient to patient. This study is distinct from previous reports because it focuses on defining transcription factors and pathways prior to onset of RA as opposed to longstanding established RA, although we also confirmed our results in validation cohorts of early and established RA^9–11^. This approach requires peripheral blood cell analysis because synovial tissue is not accessible in at-risk individuals.

To test this hypothesis, the Allen Institute for Immunology-UCSD-CU Transition to Rheumatoid Arthritis Project (ALTRA) identified at-risk individuals with elevated ACPAs, early RA patients and controls. We evaluated peripheral blood mononuclear cells (PBMCs) using single cell technologies to define the transcriptome and chromatin accessibility. Our initial recent analysis of this population using individual omics data identified broad-based evolving immune activation of PBMCs as participants progress to clinical RA^12^, but we could not address individualized mechanisms that account for the diversity of responses to targeted agents.

Here, we presented a novel, per-individual multi-omics integrative approach to determine if there is a common set of drivers that can induce aberrant immunity. We discovered a distinctive TF signature enriched in PBMCs of at-risk individuals as well as early and established RA patients. These signature TFs regulate key pathogenic processes in RA (e.g., SUMOylation, RUNX2, YAP1, NOTCH3, and β-Catenin). Unexpectedly, this signature was not confined to a single cell type but was distributed across cell types in a highly participant-specific manner. We biologically validated these predictions, demonstrating that despite the diverse cellular origins, these distinct “senders” deliver a shared core of signature inflammatory mediators to “receiver” cells *in vivo*. Furthermore, we showed that these signature mediators are actively expressed *in vivo* by RA synovial cells and that identifying an individual’s specific signature-bearing cell type is associated with clinical response to targeted therapy. Interestingly, the signature cell types can vary over time in an individual even though the inflammatory mediator profile remained relatively stable as at-risk participants progressed to clinical RA. This diversity might contribute to highly variable clinical responses to targeted therapeutics in RA patients.

## Results

### Integrative single cell analysis reveals cell types in At-Risk and CON individuals

Peripheral blood mononuclear cells (PBMCs) were obtained from 26 ACPA positive (At-Risk) and 35 age- and sex-matched controls (CON) to serve as discovery cohorts. To evaluate the persistence of discovered signatures across the clinical disease spectrum, we utilized two additional independent validation cohorts with a total of 11 patients (6 with early RA and 5 with established RA). scATAC-seq and scRNA-seq was then performed on 72 samples (**Fig. 1A, Supplementary Table S1**).

**Fig. 1.**
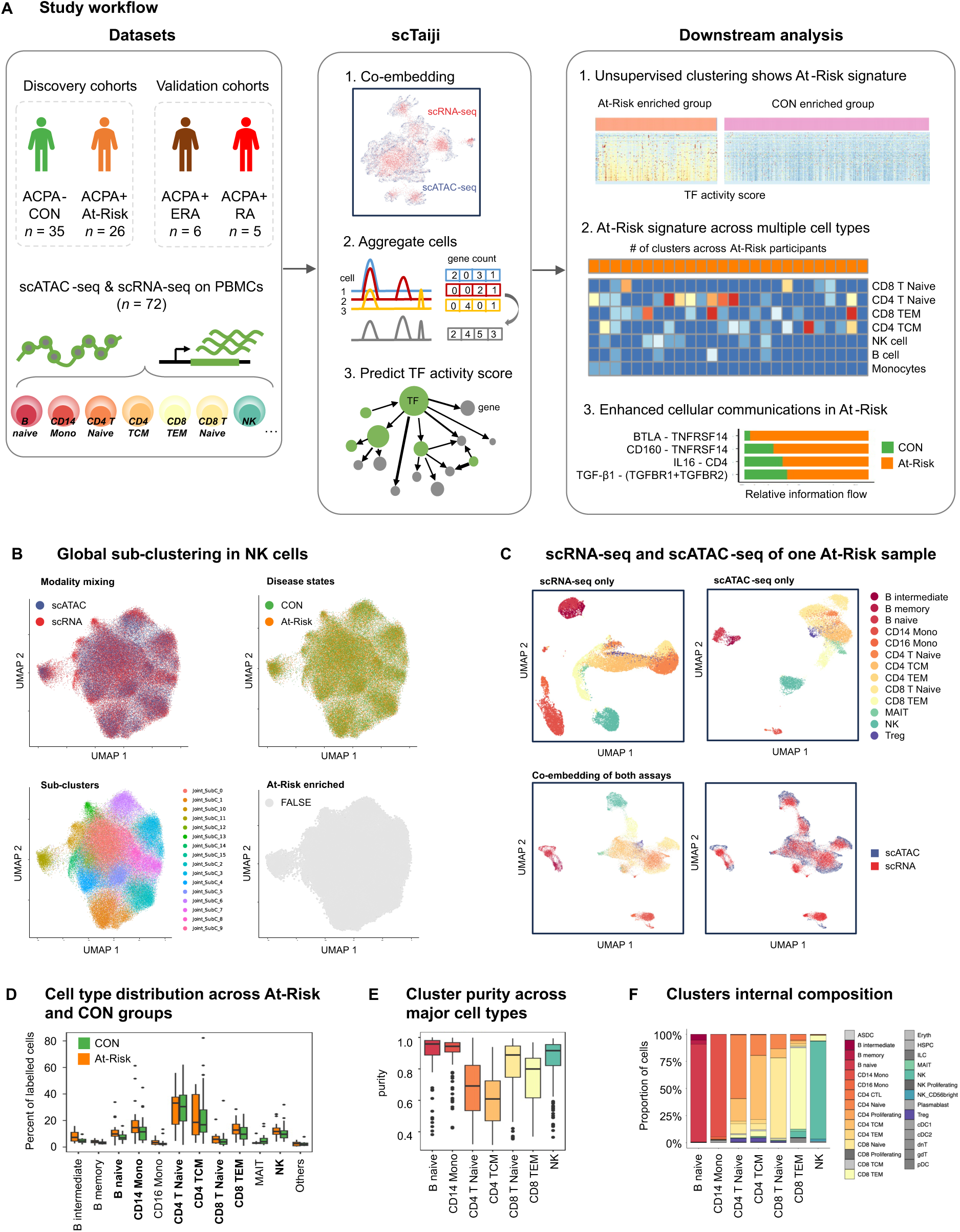
Study overview of multi-omics integrative analysis. **(A)** Schematic overview of the study design and analytical workflow. Discovery cohorts consisted of age- and sex-matched controls (CON, n = 35) and ACPA-positive individuals lacking clinical synovitis (At-Risk, n = 26). The validation cohort comprised 11 clinical RA patients (early RA [ERA], n = 6; established RA, n = 5). Paired scATAC-seq and scRNA-seq were performed on PBMCs. Following per-individual reference mapping and co-embedding, cells were aggregated into pseudo-bulk profiles. The scTaiji pipeline was utilized to predict global transcription factor (TF) activity by constructing a regulatory network per pseudo-bulk cluster. Downstream analysis identified a multi-cell-type At-Risk regulatory signature and enhanced intercellular communication networks. **(B)** Standard global integration fails to resolve disease-specific cellular states. UMAP projections of NK cells derived from a conventional pooled integration workflow (L2-normalized CCA). While the approach achieves successful modality mixing (scRNA-seq and scATAC-seq; top left) and defines multiple global sub-clusters (bottom left), At-Risk and CON cells remain highly co-mingled and indistinguishable across the integrated space (top right). Consequently, no sub-clusters are significantly enriched for the At-Risk cohort (0 clusters at FDR < 0.05 and Fold Change > 1.5; bottom right), highlighting the limitations of global clustering in capturing subtle, patient-specific pathogenic variance. **(C)** UMAP projections demonstrating high-quality multi-omics integration for a representative At-Risk sample. Panels show scRNA-seq cells alone (far-left), scATAC-seq cells alone (middle-left), and the co-embedding of both assays colored by cell type (middle-right) and assay modality (far-right). **(D)** Boxplot comparing the percentage distribution of annotated major cell types between the At-Risk (orange) and CON (green) cohorts, demonstrating similar overall cell type proportions. (E) Boxplot depicting pseudo-bulk cluster purity scores across the seven most abundant cell types analyzed in the downstream study (B naive, CD14 Mono, CD4 T Naive, CD4 TCM, CD8 T Naive, CD8 TEM, and NK cells). **(F)** Stacked bar chart illustrating the internal cellular composition of the pseudo-bulk clusters across the major cell types, confirming that lower purity in T cell compartments is primarily driven by the admixture of closely related subtypes rather than cross-lineage contamination.

Rigorous quality control was performed for both modalities (**Supplementary Fig. S1A, B**). We initially evaluated the data using a standard pooled global integration workflow (**Supplementary Fig. S1C-F**). While this conventional approach confirmed robust cell-type annotations, it could not identify patient-specific biological variance, which is crucial for understanding how heterogeneous mechanisms converge to a common phenotype in RA. Specifically, global clustering did not yield At-Risk-enriched sub-clusters; instead, At-Risk and CON cells were largely indistinguishable and co-mingled across the integrated space (demonstrated for NK cells in **Fig. 1B**, and across all other major lineages in **Supplementary Fig. S2**). This observation aligns with a recent report from the ALTRA consortium, which found that standard single-cell profiling reveals only minimum, sparsely distributed cellular differences between pre-RA and healthy individuals^12^. Notably, transcriptome-only analysis failed to distinguish the clinical cohorts (**Supplementary Notes**), demonstrating that integrating chromatin accessibility is essential to uncover subtle pathogenic signatures.

Therefore, we implemented a new per-individual, reference-based mapping approach for cell annotation, followed by scRNA-seq and scATAC-seq co-embedding. This per-participant approach offered complementary novel critical insights that would otherwise be masked by global integration (Full comparison results are provided in the Supplementary Notes). Both modalities were diffused evenly across the co-embedding space, demonstrating excellent integration across assays and samples without batch effects **(Fig. 1C**). Cells within each co-embedded cluster were aggregated to generate "pseudo-bulk" multi-omics profiles. After quality control, 1699 pseudo-bulk clusters were retained, comprising 757,250 scRNA-seq cells and 1,005,839 scATAC-seq cells from the 72 discovery and validation samples.

Our downstream analyses focused on the seven highest abundant major cell lineages in most of our analyses to assure sufficient data for analysis—CD14 monocytes (CD14 Mono), CD4 naive T cells (CD4 T Naive), central memory CD4 T cells (CD4 TCM), CD8 naive T cells (CD8 T Naive), effector memory CD8 T cells (CD8 TEM), B naive cells, and natural killer (NK) cells—which collectively accounted for >90% of all cells. The cell type distribution was similar across the At-Risk and CON groups (**Fig. 1D**). Cluster purity across these major cell types was high (mean > 0.8 for B naive, CD14 Mono, and NK clusters). While T cell subsets exhibited modestly more diverse purity scores, detailed analysis confirmed this was driven by the expected admixture of closely related T cell subtypes rather than cross-lineage contamination (**Fig. 1E, F**). Thus, one potential limitation of the analysis is that the ability to distinguish between CD4 T Naïve and TCM cells was somewhat less robust than for other cell types.

### scTaiji analysis reveals distinctive TF patterns and At-Risk signature

We applied the scTaiji pipeline to the aggregated pseudo-bulk clusters to compute PageRank scores, which represent the global regulatory influence of each transcription factor (TF) within the gene regulatory network (**Fig. 1A**). Unsupervised clustering of the TF activity matrix identified five K-means groups (G1 through G5) defined by distinct, lineage-specific TFs (**Fig. 2A, Supplementary Fig. S3D, Supplementary Table S3**). For instance, KLF4, which regulates monocyte differentiation, was G1-specific. G1 was enriched with CD14 monocytes (**Fig. 2B**). T-bet (encoded by TBX21) and EOMES displayed high activities in G3 where CD8 TEM and NK were the most abundant cell types with 37.9% and 40.3%, respectively. Those two genes are responsible for the cell fates of memory CD8+ T cells and natural killer cells^13^. Interestingly, more than half (409/640) of the TFs were G2-specific and their z scores were significantly higher in G2 compared to other groups. More than 80% of the TFs were identified as key TFs for only one Kmeans group, suggesting the Kmeans groups had unique active TF patterns (**Fig. 2C**).

**Fig. 2.**
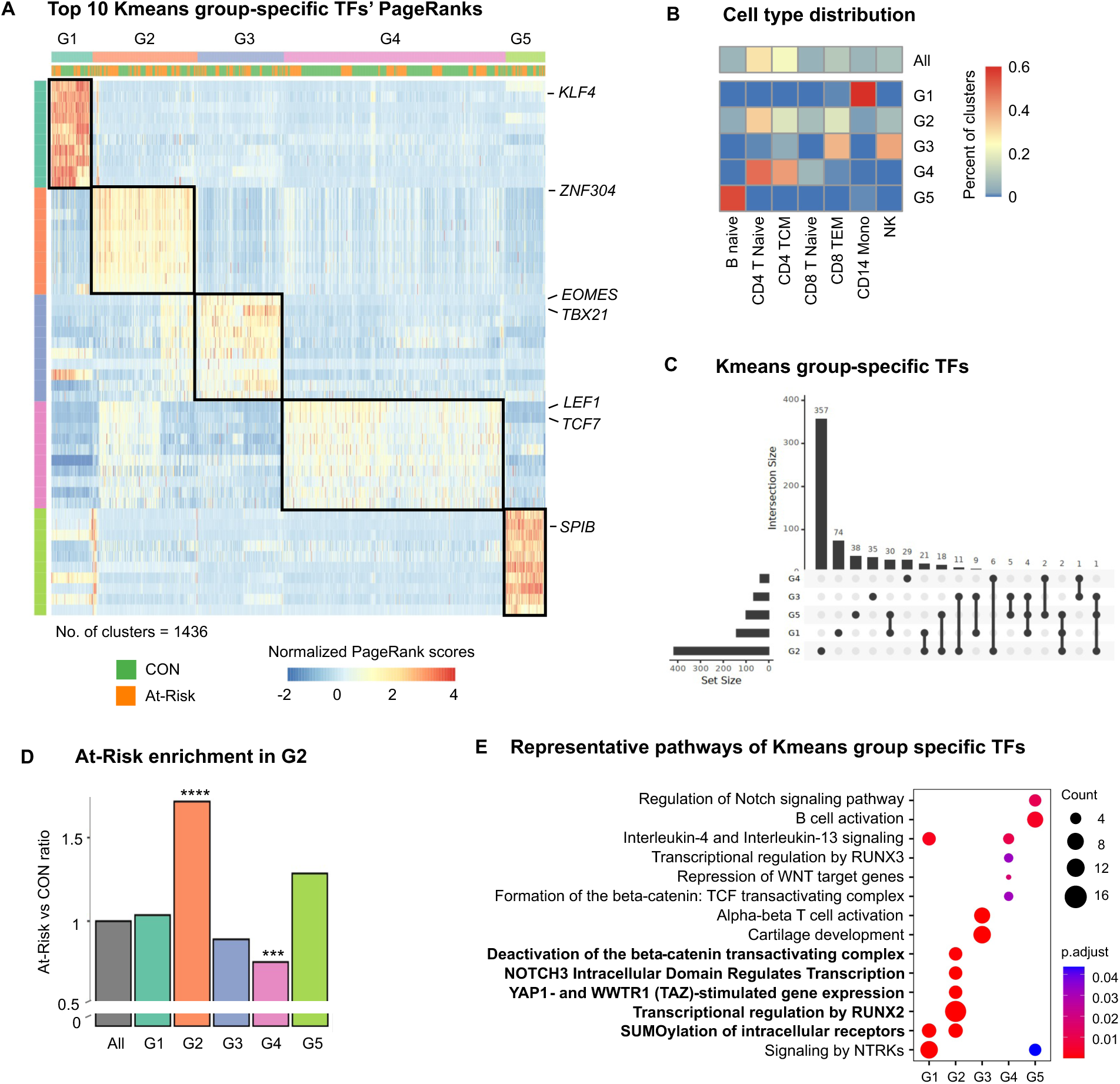
Unsupervised clustering identified signature TFs and pathways. **(A)** Heatmap displaying the normalized PageRank scores for the top 10 K-means group-specific TFs across all pseudo-bulk clusters (n = 1436). Unsupervised clustering identified five distinct TF activity groups (G1–G5). Key group-specific TFs are highlighted on the right (e.g., KLF4 for G1; ZNF304 for G2; EOMES and TBX21 for G3). **(B)** Heatmap illustrating the proportional distribution of the seven major cell types within each K-means group (G1–G5) compared to the overall dataset ("All"). While G1, G3, G4, and G5 are dominated by specific cell types, the G2 state uniquely spans all seven major compartments. **(C)** UpSet plot depicting the intersection size of group-specific TFs across the five K-means groups, highlighting that the majority of defining TFs are highly specific to a single K-means group. **(D)** Bar plot quantifying the relative ratio of At-Risk to CON clusters within each K-means group compared to the baseline dataset ratio ("All"). The multi-lineage G2 group exhibits highly significant enrichment for At-Risk clusters. **(E)** Dot plot showing representative Reactome pathways enriched among the signature TFs specific to each K-means group. Node size corresponds to the number of TFs (count), and color indicates the adjusted p-value. Key pathogenic RA pathways driving the G2 signature are highlighted in bold (e.g., Deactivation of the β-catenin transactivating complex, NOTCH3 signaling, YAP1/WWTR1 expression, RUNX2 regulation, and SUMOylation); Chi-squared test, ***p< 0.001, ****p < 0.0001.

### G2 is a multi-lineage group enriched with At-Risk and reveals an RA TF signature

The five Kmeans groups showed distinct cell type and disease state compositions (**Supplementary Table S4-5**). While G1, G3, G4, and G5 were primarily dominated by specific lineages (monocytes, CD8 TEM/NK, homeostatic CD4 T cells enriched in controls, and B cells, respectively), G2 was unique in that it was multi-lineage, encompassing all seven major cell types. (**Fig. 2B**).

Importantly, G2 was significantly enriched in At-Risk clusters compared with CON (73% higher in At-Risk vs. CON, adjusted by the null distribution, p-value < 0.0001; Chi-squared test) and G4 was modestly enriched in CON clusters (24% higher in CON, p-value < 0.001; Chi-squared test) (**Fig. 2D**). G2-specific TFs governed multiple pathways implicated in the pathogenesis of RA including *SUMOylation of Intracellular Receptors*^14^, *Transcriptional regulation by RUNX2*^15^, *YAP1 and WWTR1-stimulated Gene Expression*^16^, *NOTCH3 Intracellular Domain Regulates Transcription*^17^, and *Deactivation of the β-Catenin Transactivating Complex*^18^ Reactome pathways (**Fig. 2E; Supplementary Notes**). The TFs and the representative target genes identified by our analysis are shown in **Supplementary Table S6**. These TFs are thereafter referred to as the *RA TF signature* and their enriched pathways as *RA signature pathways*.

This "RA TF signature" was consistently expressed across multiple distinct cell lineages in the blood (**Fig. 3A**). When comparing the proportion of G2 clusters per cell type, CD4 T Naive, CD4 TCM, CD8 T Naive, and CD8 TEM showed significant enrichment in the At-Risk group compared to CON (**Fig. 3B**). These cell types displayed a common set of pathogenic signature pathways despite originating from different lineages **(Fig. 3C**), indicating systemic, multi-cellular regulatory dysregulation. Overall, the top RA signature TFs determined by unsupervised clustering showed significantly higher PageRank scores in G2 compared to other groups across all cell types (**Supplementary Fig. S3E**).

**Fig. 3.**
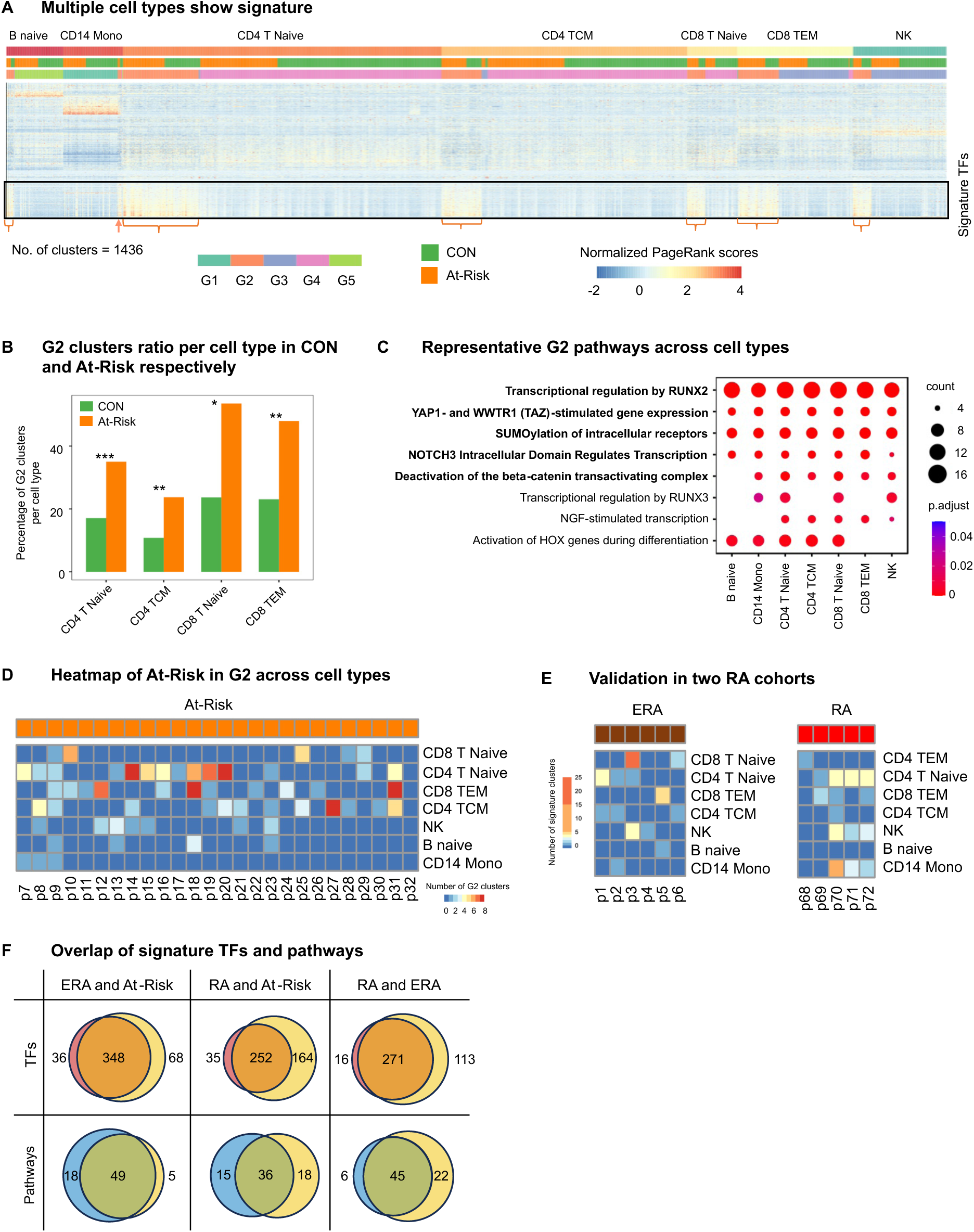
RA TF signature is broadly distributed across cell types and varies across individuals. **(A)** Heatmap of normalized PageRank scores for the signature TFs defining the pathogenic G2 state, plotted across the seven major immune cell types. Brackets and arrows highlight the consistently elevated TF activity profiles across diverse cellular compartments. **(B)** Bar plot illustrating the percentage of total G2 clusters mapped to specific T cell types, comparing the CON and At-Risk cohorts. The At-Risk group shows significant enrichment of the signature within CD4 T Naive, CD4 TCM, CD8 T Naive, and CD8 TEM populations. **(C)** Dot plot detailing the enrichment of representative G2 signature pathways across the different cell types, demonstrating that distinct cell types converge on a shared set of pathogenic regulatory programs. **(D)** Heatmap depicting the absolute number of G2 signature-bearing clusters per cell type for each individual participant in the At-Risk discovery cohort (p7 to p32). The distribution of signature cell types is highly individual-specific. **(E)** Heatmap illustrating the number of signature clusters per cell type across individuals in the RA validation cohort (early and established clinical RA). The data confirm that the multi-cell-type, patient-specific nature of the regulatory signature persists robustly after the onset of clinical disease. **(F)** Venn diagram showing the overlap of signature transcription factors (left) and signature pathways (right) between RA validation and At-Risk discovery groups; Chi-squared test, *p < 0.05, **p < 0.01, ***p <0.001.

### Patterns of cell types with the RA TF signature vary across individuals

We next evaluated which specific cell types carried this regulatory signature within individual participants. The distribution of signature-bearing cell types was highly heterogeneous across the 26 At-Risk individuals (**Fig. 3D**). Some individuals exhibited the signature broadly across nearly all major cell types (e.g. participants 8, 9, 23), while others harbored it predominantly within specific compartments, especially T cells (e.g. participants 10, 20, 27). One At-Risk individual lacked a signature cell type (p32), although we cannot rule out that this occurred in a less common cell type that was beyond the resolution of this analysis.

Different cell types also displayed diverse distribution patterns across individuals. CD4 Naive and/or CD4 TCM had wider appearances in many participants while B naive and CD14 monocytes were only found in a few. Some CON participants also displayed these signatures although the number of clusters was significantly less than At-Risk, particularly for certain T cell subsets (p-value < 0.005; Wilcoxon rank-sum test) (**Supplementary Fig. S4A**).

#### Validation in two RA cohorts

To confirm that this pre-clinical signature persists after clinical onset, we applied our analytical framework to the clinical validation cohorts. The RA TF signature was robustly detected across all 11 RA patients in the two validation cohorts whether they were analyzed as a group or separately (early RA and established RA), confirming its stability as a pathogenic module regardless of clinical duration or treatment (**Fig. 3E, F** show separate analyses for each cohort and **Supplementary Fig. S4B-D** show combined analysis). The findings were consistent across RA groups despite differences in disease duration and treatment. As with the At-Risk cohort, the distribution of signature-bearing cell types was diverse in the validation cohorts.

### Enhanced cellular communication networks in At-Risk individuals

To understand how these signature cells transmit inflammatory signals, we analyzed cell-cell communication (CCC) networks. The At-Risk cohort displayed a significantly higher number of differential interactions and greater interaction strength within signature clusters compared with CON (**Fig. 4A**). Cellular communications between all signature cell pairs were more pronounced in At-Risk group except partial interactions where CD8 T cells are involved (**Fig. 4B; Supplementary Fig. S4E**). It is worth noting that the number and intensity of the total CCC aggregating all the clusters from all the Kmeans groups were comparable between the At-Risk and CON groups (**Supplementary Fig. S4F**). This lack of global divergence indicates that pathogenic signaling is specifically localized within the G2 signature clusters; consequently, restricting the analysis to this subset is essential to isolate the disease-specific signals that would otherwise be masked by baseline homeostatic communication. Note that we cannot determine where this interaction occurs, such as synovium, lymphoid tissue, or mucosa. Because predicted sender mediators are elevated in serum (see below), it could also occur in blood without direct cell-cell contact.

**Fig. 4.**
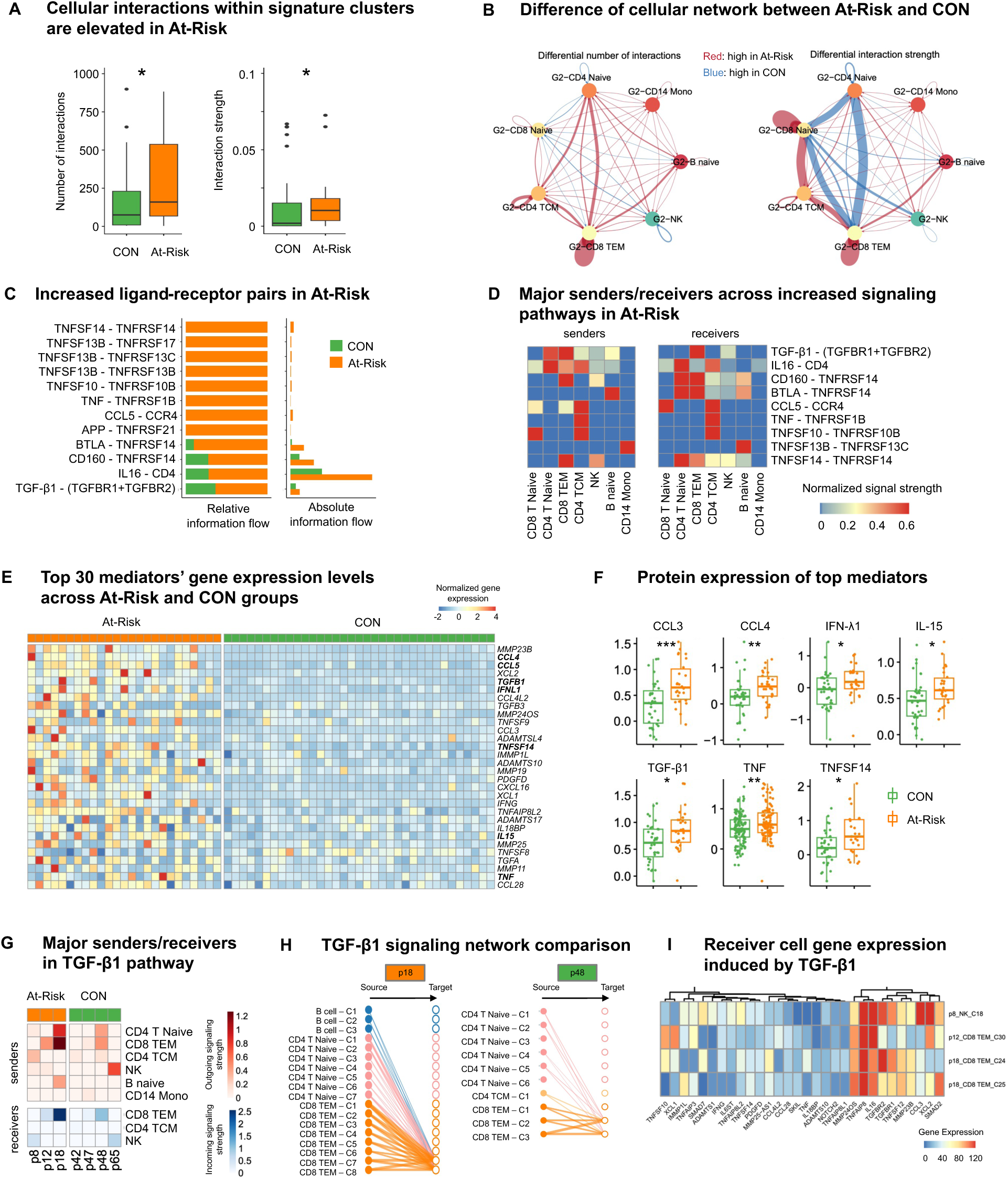
Enhanced cell-cell communication networks and universal pathogenic mediators define the At-Risk state. **(A)** Boxplots demonstrating a significantly increased number of differential cell-cell interactions (left) and interaction strength (right) within the signature clusters of the At-Risk cohort compared to CON. **(B)** Network diagrams illustrating the differential number of interactions (left) and interaction strength (right) directly between the seven major signature cell types in the At-Risk cohort. Line thickness indicates the magnitude of signaling elevation. Red edge indicates more (left) / stronger (right) interactions in At-Risk and blue is vice versa. **(C)** Bar charts ranking the top elevated ligand-receptor pairs in the At-Risk group based on relative information flow (left) and absolute information flow (right). The total information flow is calculated by summing the probability of all communications between the signature clusters. **(D)** Heatmap displaying the normalized signal strength of the major sender (left) and receiver (right) cell types orchestrating the top elevated signaling pathways in At-Risk individuals, highlighting multi-cellular cross-talk. **(E)** Heatmap showing the normalized gene expression of the top 30 pathogenic mediators—identified via a random forest classification model—across the At-Risk and CON cohorts. Top 30 predictor cytokines are uniformly more active in At-Risks compared to controls. Example genes include *MMP23B*, *CCL4*, *IL12A*, *TNFSF14*, *IL15*, *NOTCH1*, *CCL5*, and *TGFB1.* **(F)** Boxplots biologically validating the transcriptomic model, showing significantly elevated circulating protein levels of key predicted mediators (CCL3, CCL4, IFN-λ1, IL-15, TGF-β1, TNF, and TNFSF14) in At-Risk patient serum compared to CON. **(G)** Heatmap detailing the heterogeneous major sender and receiver cell types utilized for the TGF-β1 signaling pathway across At-Risk and CON individuals. (**H**) Network diagrams directly comparing the complexity and density of the TGF-β1 signaling networks within signature clusters between a representative At-Risk (p18) and CON (p48) individual. **(I)** Heatmap confirming the active downstream biological consequence of this signaling, demonstrating the robust expression of key receiver genes (e.g., *TGFBR1* and *TGFBR2*) induced by TGF-β1 in targeted receiver cells; Wilcoxon rank-sum test, *p < 0.05, **p < 0.01, ***p < 0.001.

Analyzing specific ligand-receptor pairs revealed elevated signaling flow in the At-Risk group for key inflammatory networks, notably including IL16 - CD4, CD160 - TNFRSF14, TGF-β1 - (TGFBR1+TGFBR2), and BTLA - TNFRSF14 (**Fig. 4C**). The major senders and receivers orchestrating these enriched pathways spanned multiple distinct cell types, further illustrating the multi-cellular nature of the pre-clinical state (**Fig. 4D**).

### Classification model identifies universal pathogenic mediators

To identify the most robust systemic drivers of the At-Risk state, we developed a random forest classification model trained exclusively on the At-Risk and CON discovery cohorts. Rather than relying on highly abundant cell types, the model utilized the maximum gene expression of candidate mediators per participant, ensuring that all cells were equally weighted. The model successfully distinguished At-Risk from CON individuals (**Supplementary Fig. S4G**), identifying a core set of predictor genes (**Supplementary Fig. S4H**).

Pathogenic mediators such as *MMP23B*, *CCL4*, *CCL5*, *TGFB1*, and *IFNL1* were significantly higher across the At-Risk cohort compared with CON (**Fig. 4E**). *MMP23B*, which emerged as a top predictor in classification model, regulates the Kv1.3 potassium channel, which has been implicated in autoimmunity^19^. Although a common set of pathogenic genes were shared across At-Risk participants, the cell types and individual mediators that were most likely to produce the specific gene were variable (**Supplementary Fig. S4I**). The top 30 predicted mediators are referred to as *signature genes*.

### Biologic validation of computational predictions

#### Biologic validation of signature gene predictions

Functional validation of our transcriptomic predictions was initially performed by measuring serum protein expression levels of key top signature gene predictors. Concordant with our genomic model, circulating protein levels of CCL3, CCL4, IFN-λ1, IL-15, TGF-β1, TNF, and TNFSF14 were significantly elevated in At-Risk group (**Fig. 4F**).

#### Biological validation of communication networks

As an example of a robust communication network gene, we examined the TGF-β1 signaling network. The major senders and receivers of TGF-β1 signals varied across individuals but converged on strong pathogenic communication (**Fig. 4G**). For instance, contrasting a representative At-Risk individual (participant p18) with a healthy control (participant p48) vividly illustrated the dense, multi-lineage signaling network unique to the disease state (**Fig. 4H**). Furthermore, we confirmed the functional biological consequence of these signals by verifying the significant induction of downstream TGF-β1 - responsive genes in receiver genes in the targeted receiver cells (CD8 TEM) such as TGFBR1, TGFBR2, and TNFAIP8 (**Fig. 4I**).

We also validated the predicted receiver cell response for other key signaling networks. For instance, IL16 - CD4 signaling pathway, which has been implicated in RA^20^, showed significantly stronger signals in At-Risk group (**Supplementary Fig. S5A-B**). We observed significantly elevated expression of key downstream target genes, including *CDKN1B*, *IL32*, and *TNFAIP8*, in the predicted receiver cell types (**Supplementary Fig. S5C**).

Two other pathways that are elevated in At-Risk are CD160 and BTLA signaling pathways (**Supplementary Fig. S5D, G**). While NK cells were the most common senders for CD160 signaling, B cells acted as the exclusive senders for BTLA signaling; both pathways targeting a diverse range of receiver cell types (**Supplementary Fig. S5E, H**). Like TGF-β1, we confirmed significantly elevated expression of predicted key downstream genes induced by the sender mediator in the predicted receiver cell types, including *TNFRSF14* and *TNFAIP8* (**Supplementary Fig. S5F, I**).

#### Biological validation of signature genes in RA synovium

To determine whether the blood-derived systemic signature reflects joint pathology, we evaluated synovial scRNA-seq data from the independent Accelerating Medicines Partnership (AMP) cohort^11^. The top circulating predictor genes identified in our blood cohorts were highly expressed in the rheumatoid synovium. As observed in the pre-clinical PBMCs, the specific cell types expressing these signature mediators were highly variable across individual RA patients (**Fig. 5A**). Furthermore, by explicitly incorporating tissue-resident stromal populations into our analysis, we discovered that synovial fibroblasts also act as major local sources for a distinct module of the systemic signature gene predictors and can also serve as receivers. Granular details regarding the specific expression mapping across all finely resolved synovial sub-clusters are provided in the Supplementary Notes and **Supplementary Fig. S6A, B**.

**Fig. 5.**
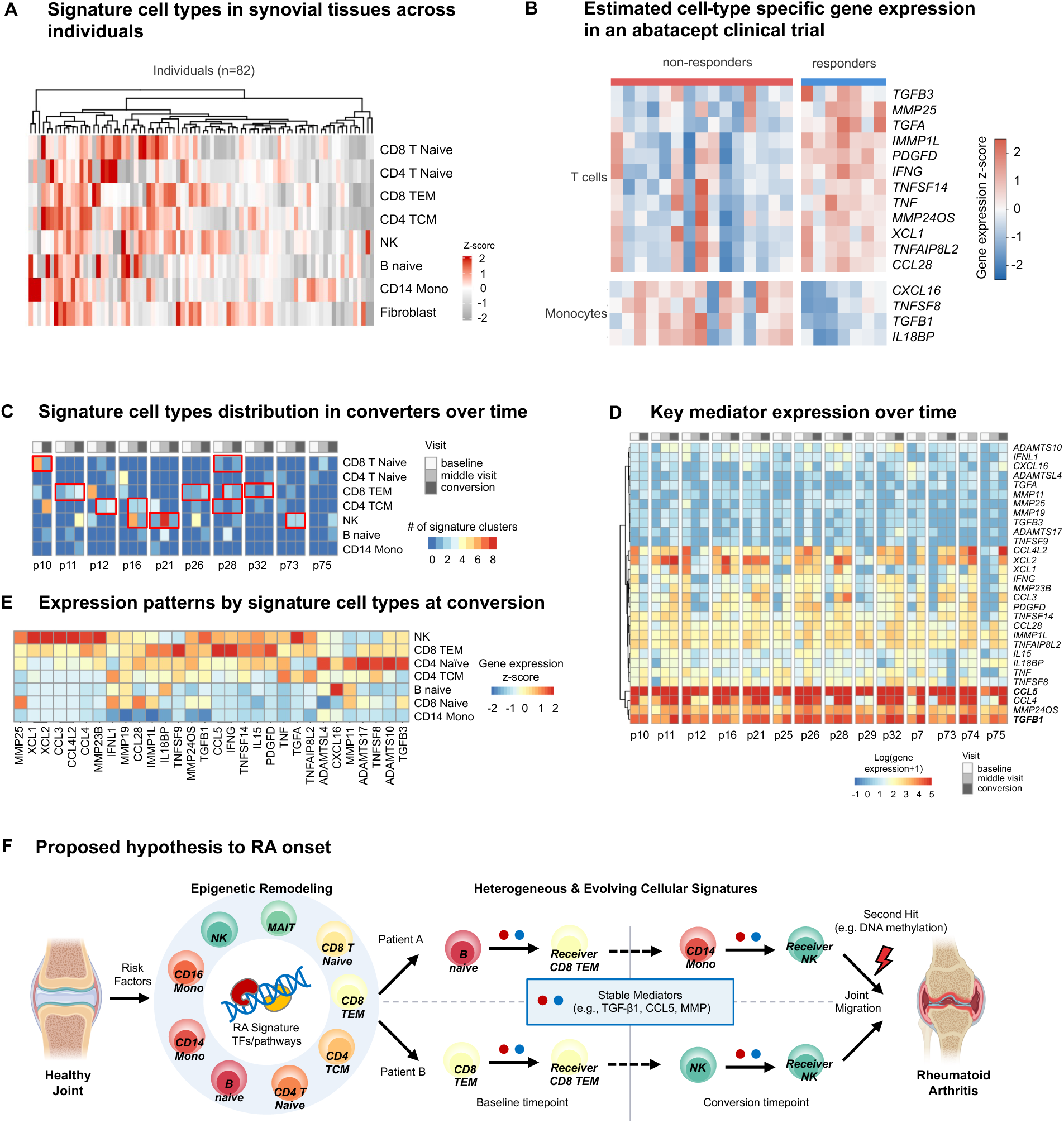
Biologic validation in target tissues and longitudinal dynamics of the RA signature. **(A)** Heatmap depicting the scaled pseudo-bulk gene expression (z-score) of top blood-derived signature mediators across single-cell populations in 82 inflamed synovial tissue samples (AMP cohort). This analysis confirms robust tissue-level expression and highlights tissue-resident synovial fibroblasts alongside infiltrating immune cells as major patient-specific sources of pathogenic mediators. **(B)** Heatmap of imputed cell-type-specific gene expression derived from the bulk PBMC RNA-seq deconvolution of an abatacept clinical trial. Patients are stratified into non-responders and responders, revealing that therapeutic failure is associated with strong signature gene expression hosted predominantly within the monocyte compartment. **(C)** Heatmap illustrating the longitudinal tracking of signature-bearing cell types in At-Risk individuals who converted to clinical RA where conversion data was available (baseline, middle visit, conversion). The specific cellular hosts of the signature exhibit significant plasticity and dynamic evolution over time. The red boxes show participants where the conversion cell type recapitulated a previous cell type (p = 0.0272). **(D)** Heatmap tracking the log-normalized expression of the core mediator genes within these same converting individuals over time. Despite the shifting cellular hosts, the inflammatory mediator profile remains remarkably stable from the pre-clinical phase through conversion. **(E)** Heatmap detailing the specific gene expression output contributed by each signature cell type at the precise time of clinical conversion. **(F)** Schematic model of the proposed hypothesis for RA onset. Genetic and environmental risk factors might trigger epigenetic remodeling, establishing the RA TF signature within evolving immune cell types. Examples of potential senders and receivers are shown, although there are many other potential combinations. Despite their diversity, these cellular hosts converge to deliver a stable core of inflammatory mediators (e.g., TGF-β1, CCL5) that activate receiver cells including tissue-resident fibroblasts. There are multiple places where targeted interventions might be beneficial, including the sender cell, the sender mediator, or the receiver cell.

#### Biological validation of therapeutic relevance

We then evaluated bulk PBMC RNA-seq data from a published clinical trial of abatacept to test the clinical relevance of the signature’s cellular origin^21^. Deconvolution analysis revealed that patients who did not respond to abatacept (non-responders) exhibited strong signature gene expression primarily within monocytes, whereas responders were characterized by T cell-driven signatures (**Fig. 5B**). This observation is consistent with the putative mechanism of action for co-stimulation inhibition with blockade of T cell activation in RA. In other words, only individuals with a signature in T cells responded to a T cell directed therapy.

### Longitudinal analysis: Evolving signature cell types and stable mediator expression

#### Evolving signature cell types in converters

Longitudinal tracking of At-Risk individuals who converted to clinical RA revealed that the specific cell types carrying the RA signature could evolve over time (**Fig. 5C**). Importantly, the specific cell types carrying the RA signature typically returns to a previous cell type when participants transition to RA (p = 0.0272).

#### Stable mediator profiles in converters

Despite signature cell type variability, the core inflammatory profile remained surprisingly stable. Key mediator genes, such as *CCL5* and *TGFB1*, were stably expressed from baseline through conversion, regardless of which signature sender cell type was implicated (**Fig. 5D**). At the time of clinical transition, specific cell types assumed specialized roles in producing this mediator profile (**Fig. 5E**). For instance, NK cells are highly active producers of chemokines like *XCL1* and *XCL2* while CD8 TEM cells make the greatest contribution to *CCL5* and *TNFSF8* expression. We also observed a prominent activation signature in CD4 Naive T cells, consistent with our previous observation^12^. Together, these longitudinal data support a model (**Fig. 5F**) where diverse and evolving immune cell types producing common inflammatory mediator modules converge to deliver a stable, unified pathogenic signal that drives the transition to clinical rheumatoid arthritis.

## Discussion

Our study identifies a conserved, pathogenic TF signature present in at-risk ACPA+ individuals and patients with RA. The signature TFs orchestrate key pathways implicated in RA pathogenesis like *SUMOylation*, *RUNX2*, *YAP1*, *NOTCH3*, and *β-Catenin* pathways^14–18^. Strikingly, the unifying signature is not confined to a single cell type but is hosted by a diverse and plastic array of immune cell types in a patient-specific manner. This observation helps explain how heterogeneous cellular drivers can produce a common clinical phenotype. We also demonstrate that these varied cellular hosts converge to produce a stable, shared profile of inflammatory mediators. The clinical relevance of this model was validated in two separate RA cohorts with varying disease duration and treatment, and a clinical trial confirming that the findings are robust, durable and functionally important. Our observations offer a new paradigm for RA pathogenesis that mechanistically explains therapeutic variability and could provide a framework for personalized medicine.

Our computational predictions were biologically validated across multiple modalities. First, key inflammatory mediators that we identified (e.g., TGF-β1, CCL4, IL-15, TNF) were confirmed to be elevated at the protein level in patient serum. This observation suggests that communication between senders and receivers could occur in blood rather than requiring close apposition. Second, our predicted cell-cell communication networks were functionally validated by confirming that the receiver cell types express genes *in vivo* that are regulated by their paired sender cell signal. Third, synovial tissue from the AMP cohort revealed that the blood-derived predictors are robustly expressed in inflamed synovium. Importantly, incorporating tissue-resident stromal populations showed that synovial fibroblasts act as major local sources and receivers for a distinct module of these mediators, directly bridging systemic immune dysregulation and local joint inflammation. Finally, deconvolution of an abatacept clinical trial supported our therapeutic hypothesis: a T cell-targeted agent was not effective when the patient’s pathogenic signature was primarily present in monocytes rather than T cells.

One of the most intriguing observations is the individualized and plastic nature of this RA signature. Each patient exhibited a distinct combination of signature-bearing cell types, suggesting a stochastic component to remodeling the epigenome. Longitudinal tracking revealed that the specific immune cell types carrying this signature could evolve somewhat over time. However, the cell type usually returns to an earlier lineage at the time of conversion and the core inflammatory mediator profile remains remarkably stable despite being produced by different senders. This model—where diverse and fluctuating cellular drivers converge on a consistent inflammatory output—provides a mechanistic explanation for the highly variable clinical responses observed with targeted, cell-centric therapies. It suggests that the responding cells are agnostic to the sender’s identity as long as the pathogenic mediators are delivered to a receiver.

Our methodological approach distinguishes this work from previous pre-RA studies that predominantly relied on the transcriptome-only data from established RA synovium or peripheral blood. The integration of transcriptome and chromatin accessibility data revealed patient-specific, multi-lineage pathways that would have been missed by transcriptome-only analysis^22^ (**Supplementary Notes**). Previous single-omics analyses primarily highlighted singular pathogenic cell types such as CD4+ T naïve cells or CCR2+ CD4+ T cells^12,23^. However, preponderance of a single pathogenic cell type would not explain why targeted T cell agents like abatacept failed in a significant subset of patients^24^. Our integrated analysis demonstrates that B cells, CD8+ T cells, monocytes and NK cells also exhibit the signature and produce the same pathogenic mediators as CD4+ T cells in some participants.

Signature cell type plasticity over time was surprising and likely reflects stochastic events leading to immune activation, perhaps due to repeated stimulation at mucosal surfaces by circulating cells. The observed cellular remodeling offers a mechanistic explanation of longitudinal enhancement of serum protein mediator production during RA development^25^. Because the underlying TF signature remains consistent across the at-risk, early and established RA phases, it likely represents a “primed” molecular state. A second “hit”, such as DNA methylation^26^, might push someone from “at-risk” to clinical autoimmunity. While it is tempting to target an individual pathogenic cell type, this should be tempered by the observation that it can vary over time, and many individuals have multiple signature cell types. Interestingly, the inflammatory mediator profile is more stable and might be more useful as a biomarker for stratification and developing personalized approaches to treatment.

The surprising overlap of the RA TF signature and pathways across multiple cell types suggests that a shared upstream mechanism shaping RA epigenome. Environmental and mucosal stresses, especially in the airway due to its critical role in the RA, are possible influences because all circulating cell types can be exposed to irritants at these sites. For example, cigarette smoke, a major RA risk factor, induces stress throughout the airway and is associated with epigenetic alterations in peripheral blood cells^27^. We also previously described DNA methylation abnormalities in circulating B cells and CD4 T cells in the at-risk population^28^, which supports this concept. It is also possible that multiple signature cell types are influenced by similar inflammatory signals, but the impact could be divergent depending on where they are imprinted (e.g., gut, lung, or synovium).

While we observed these conserved patterns across pre-clinical and established RA spectrum, we do not know whether the signature extends to other immune-mediated diseases. Recent studies have indicated the protein-level specificity distinguishing RA from Sjögren’s syndrome, SLE, and systemic sclerosis^25^, but it is possible that this signature represents a general phenomenon of autoimmunity. If so, the ultimate manifestation of a particular autoimmune disease might be determined by other factors, such as genetic predisposition and environmental influences. Nevertheless, in certain diseases like psoriasis, which responds uniformly well to Th17-directed therapies^29^, the pathogenic cellular repertoire appears much narrower than in RA. Thus, the specificity and cellular distribution of this systemic signature warrant further cross-disease investigation.

In conclusion, our study defines a distinctive, multi-cellular RA TF signature that drives an inflammatory program that spans from the pre-clinical phase through established disease. We demonstrated that diverse immune cell types can converge to deliver a shared, stable pro-inflammatory signal that could drive transition to clinical arthritis. This individualized, multi-lineage paradigm provides a mechanistic explanation for why clinical responses to targeted therapies are so highly variable in RA and establishes a foundational framework for developing personalized prognostic tests and therapeutic strategies.

## Materials and Methods

### Clinical cohorts

Four groups of participants were recruited for this study. The demographics and baseline characteristics are provided in **Supplementary Table S1**. The study design was structured as a primary discovery phase and subsequent validation phase. The discovery cohorts consisted of two groups: (1) an “At-Risk” cohort (n=26) comprising individuals positive for anti-citrullinated protein antibodies (ACPA; >2x the upper limit of normal^2^ using the anti-CCP3 IgG ELISA, Werfen, San Diego, CA) but entirely lacking clinical synovitis (joint swelling) upon rheumatological examination; and (2) age- and sex-matched control participants (CON, n=35). The 26 At-Risk individuals were selected from a larger parent cohort (n=45) strictly based on the availability of high-quality, paired scRNA-seq and scATAC-seq data from the same blood draw and enriched for individuals that converted to clinical RA. During longitudinal follow-up, 16 of these 26 At-Risk individuals converted to clinical RA based on clinical evaluation and a finding on physical examination by a rheumatologist or trained study nurse of ≥1 swollen joint consistent with inflammatory arthritis. The validation cohorts consisted of patients with a confirmed clinical diagnosis of RA to test the persistence of identified signatures across the disease spectrum. This included: (3) early RA patients (ERA, n=6) diagnosed <1 year from study enrollment based on the ACR/EULAR 2010 RA classification criteria^30^, and (4) established RA patients (n=5) identified at the time of arthroplasty with disease duration greater than 1 year, treated with various anti-rheumatic agents agents (e.g., JAK inhibitor, TNF blockers, and hydroxychloroquine). Participants were recruited at the University of Colorado Anschutz, UC San Diego, and the Benaroya Research Institute. The studies were approved by ethical review boards at all institutions, and all participants gave informed consent. Available clinical information is summarized in **Supplementary Table S1**.

### Sample preparation, scRNA-seq, and scATAC-seq

Blood was collected, and PBMCs were isolated using Leucosep tubes with Ficoll Premium and cryopreserved. For scRNA-seq, libraries were generated using a modified 10x Genomics Chromium 3’ single-cell gene expression assay with Cell Hashing as previously described^31^. For scATAC-seq, dead cells, debris, and neutrophils were excluded via FACS sorting (CD45+ viable gating). Permeabilized-cell scATAC-seq libraries were prepared using the Chromium Single Cell ATAC Gel Beads v1.1 protocol. Final libraries were sequenced on the Illumina NovaSeq platform. Detailed protocols are provided in the **Supplementary Methods**.

### Single-cell data processing and per-patient cell annotation

To preserve patient-specific biological variance, we employed a per-individual, reference-based mapping approach for cell annotation.

#### scRNA-seq data

Following alignment (CellRanger v3.1.0) and hashtag processing, rigorous quality control was performed per sample. High-quality cells were retained based on: <10% mitochondrial reads, 200–5000 detected genes, and library complexity (log10GenesPerUMI) >0.80. Doublets were removed using scDblFinder. For normalization, Seurat’s SCTransform (v2 regularized negative binomial regression) was applied to each sample individually. Each patient’s dataset was then independently projected onto the top 30 Supervised Principal Components of a validated, multimodal healthy PBMC reference^32^. QC summary plots along with statistics can be found in **Supplementary Fig. S1A** and **Supplementary Table S2**.

#### scATAC-seq data

Following CellRanger-ATAC (v1.1.0) alignment, cells were retained if they exhibited between 1000-100,000 unique fragments, 10-2000 fragment size, >50% fragments in valid regions, >20% of fragments in transcription starting site (TSS), >4 TSS enrichment score. ArchR v1.0.2 was used to generate Arrow files, doublet filtering (filterRatio=0.5), dimensionality reduction with iterative latent semantic indexing (LSI) (iterations=4), and clustering (resolution=3). QC summary plots along with statistics can be found in **Supplementary Fig.** S1C **and** Supplementary Table S2.

#### Aggregated Visualization

While all primary downstream statistical analyses were performed at the per-patient level, reference-projected embeddings from all individuals were concatenated solely to generate global UMAPs for visual overview (**Fig. S1**).

### scRNA-seq and scATAC-seq co-embedding and pseudo-bulk aggregation

scATAC-seq data were integrated with the corresponding scRNA-seq using the “addGeneIntegrationMatrix” function in ArchR with default parameters. Each cell in the scATAC-seq space was assigned a gene expression signature from its most similar cell in the scRNA-seq neighbor. Cells from both modalities were clustered in the co-embedding space in an unsupervised way. Single cells within the same cluster were then aggregated into a “pseudo-bulk” profile by summing scRNA-seq counts and combining scATAC-seq fragments. Pseudo-bulk cluster annotation was taken from the cell type occurring most frequently in the cluster. Only pseudo-bulk clusters with >2000 open chromatin peaks, >20 scATAC-seq cells and >20 scRNA-seq cells were kept for the downstream regulatory network construction.

### TF regulatory networks construction (scTaiji) and Unsupervised Clustering

For each pseudo-bulk cluster, Taiji v1.1.0 was used to construct gene regulatory networks. PageRank scores representing global TF influence were calculated based on open chromatin peak intensity, TF expression, and motif binding affinity. To identify groups of samples with similar active TF patterns, we performed PCA on the TF PageRank score matrix. Sensitivity analyses determined that retaining the top 5 principal components (PCs) optimally captured robust cell-type identities while eliminating diffuse, high-dimensional noise. Using these top 5 PCs, we performed unsupervised clustering.

We benchmarked multiple algorithms (Kmeans, hierarchical, and spectral) and determined Kmeans yielded the best clustering quality. For each sample *i*, let *a*(*i*) be the mean distance to other samples and *b*(*i*) the mean distance to samples in the nearest neighboring cluster. The silhouette coefficient is 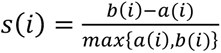. The optimal number of groups and the use of the clustering algorithm were established via Silhouette analysis (**Supplementary Fig. S3A-C**).

### Identification of Kmeans group-specific TFs and regulatees

To identify Kmeans group-specific TFs, clusters were divided into a target group and a background group. Because Shapiro-Wilk testing revealed 95% of PageRank scores did not follow normal distribution, the Mann-Whitney U Test was utilized. Specific TFs were called using double cutoffs: P-value ≤ 0.01 and log2 fold change ≥ 0.5. Results were summarized in **Supplementary Table S3**. Top regulatees were ranked by mean Taiji edge weight across signature TFs. Representative regulatees were summarized in **Supplementary Table S6**.

### Cell-cell communication analysis

CellChat (v2.1.2) was used to analyze intercellular interactions. Normalized scRNA-seq data (TPM, log-transformed) and co-embedding cell labels were used as input. Validated molecular interactions from CellChatDB v2 (excluding non-protein signaling) were used to generate individual-level communication networks, which were subsequently aggregated for group-wise comparison.

### Identification of candidate pathogenic genes related to signature group G2

We first curated a customized list of 187 genes including all the available cytokines, chemokines, growth factors, NOTCHs, MMPs, and ADMATS with gene expression in this study. The full gene list is shown in **Supplementary Table S7**. For each gene, the maximum gene expression across clusters was taken within each Kmeans group and each individual as input. Then, we identified the universal G2-important genes with mean gene expression across all patients ranked as top 50% and coefficients of variation (CV) less than 2. In total, 63 genes were identified as candidate predictors for the following classification model.

### Classification model construction

To distinguish At-Risk individuals from Controls without confounding from clinical disease, the random forest model was trained exclusively on the discovery cohort (At-Risk, n=26; CON, n=35). To mitigate cell-type abundance bias—where highly abundant T cell populations mask signals from rarer populations like B cells or monocytes—our feature engineering utilized the maximum gene expression value for each candidate mediator across the signature G2 clusters per participant (or G4 clusters for controls). This ensured potent signals from rare cells were weighted equally to those from abundant cells. The samples were split into train/test sets (7:3). Feature importance was evaluated using the recursive elimination algorithm implemented (rfe in the R package caret). Models were trained using 10-fold cross-validation (repeated 5 times) and validated on the unseen test set across 20 different random seeds.

### Synovial tissue analysis (AMP study comparison)

To confirm the expression of blood-derived predictors in target joint tissue, we analyzed the AMP Phase 2 synovial dataset^11^. Raw UMI counts were collapsed into pseudo-bulk matrices per sample and cluster. Crucially, our analysis was expanded across 77 finely resolved AMP clusters to explicitly incorporate tissue-resident stromal populations, including distinct synovial fibroblast and endothelial cell sub-populations. Normalized counts per million were averaged and visualized to assess the multi-cellular spatial distribution of the identified predictors.

### Abatacept treatment response deconvolution analysis

To assess the clinical relevance of our identified cell-specific signatures, we analyzed another cohort that participated in an independent abatacept clinical trial in RA with available RNA-seq data in public databases^21^. We utilized pre-treatment (baseline) bulk PBMC RNA-seq from the 15 non-responders and 7 responders with available data. Because abatacept targets T-cell costimulation, we deconvoluted the bulk data to isolate cell-type specific gene expression. CIBERSORT-style linear support vector regression was applied with 1,000 permutations using a PBMC single-cell RNA-seq reference^33^ (17,878 genes across monocytes, CD8⁺ T, CD4⁺ T, NK, and B cells). Following the approach of Avila Cobos and colleagues^34^, we imputed cell-type-specific expression matrices (CPM-equivalent units) by combining the bulk signal with the estimated cell-type fractions. Imputed expression values for candidate predictors were compared between non-responders and responders within each cell compartment using a two-sided Mann–Whitney U test. with Benjamini-Hochberg FDR correction. Genes with nominal p ≤ 0.05 were considered statistically noteworthy and shown in the heatmap.

### Longitudinal sample analysis

For longitudinal data, optimal K-means clustering (K=4) was established using Silhouette analysis. Multi-lineage groups were analyzed for dynamic TF activity patterns over time, and pathway overlap between cross-sectional and longitudinal datasets was evaluated.

## Data availability

scRNA-seq and scATAC-seq data from this paper are deposited in the GEO database (GSE278746). The output of this study (TF activity heatmap, individual UMAP and cellular network plots) will be available at our Taiji-altra portal (https://wangweilab.shinyapps.io/Taiji_Altra/). Additionally, we implemented an interactive interface for natural language queries of our datasets and related literature (https://huggingface.co/spaces/taijichat/Taiji_ALTRA). All other raw data are available from the corresponding authors upon request.

## Code availability

The code to reproduce the data analysis and related figures in this study can be found at https://github.com/Wang-lab-UCSD/Taiji_ALTRA

## Acknowledgments

This project was supported by grants from the Allen Institute for Immunology (GSF) and NIH (R01AR065466 to WW and GSF). We thank the study participants for their valuable time and contributions to this study. We thank the clinical research team at University of California, San Diego, University of Colorado, Benaroya Research Institute for recruitment and sample preparation. This research was supported by the Allen Institute, founded by Jody Allen – chair and co-founder of Allen Family Philanthropies, and the late Paul G. Allen – investor, philanthropist, and co-founder of Microsoft. We gratefully acknowledge their vision and generosity, which make this work possible. We thank Adam Savage for support and critical review of the manuscript, the Allen Institute, Immunology operations team for maintaining the productive research environment, and the Human Immune System Explorer (HISE) software development team for their support and dedication. This paper and the research behind it would not have been possible without HISE, a collaborative computational data analysis environment for life sciences research. We thank Christy Adams, Amelia Dayton, Aishwarya Gogate, and Upaasana Krishnan for GEO and dbGaP submssions.

## Author contributions

GSF, WW, KDD, VMH, JHB and TFB conceived and designed the project.

KN, VT, LL, AO, AW, MF, CS, JHB, CS identified and worked with the research subjects who participated and managed the project, with assistance from MLF, MKD, KAK, FZ, LKM, MC, BH, MS.

DB developed methodology and DB supervised the sample collection and processing.

PG, MW, VH, JR performed studies that generated data for the project. LO developed methodology and performed analysis. MAG, PS supervised data acquisition. LB is in charge of project management and TFB for cohort conceptualization.

CL and WW performed bioinformatics analysis with assistance from EBP and PW.

CL, WW, and GSF interpreted analytical results.

CL, WW and GSF drafted the initial manuscript.

All authors reviewed and edited the manuscript. All authors approved the final manuscript.

## Supplementary Figures

**Fig. S1.**
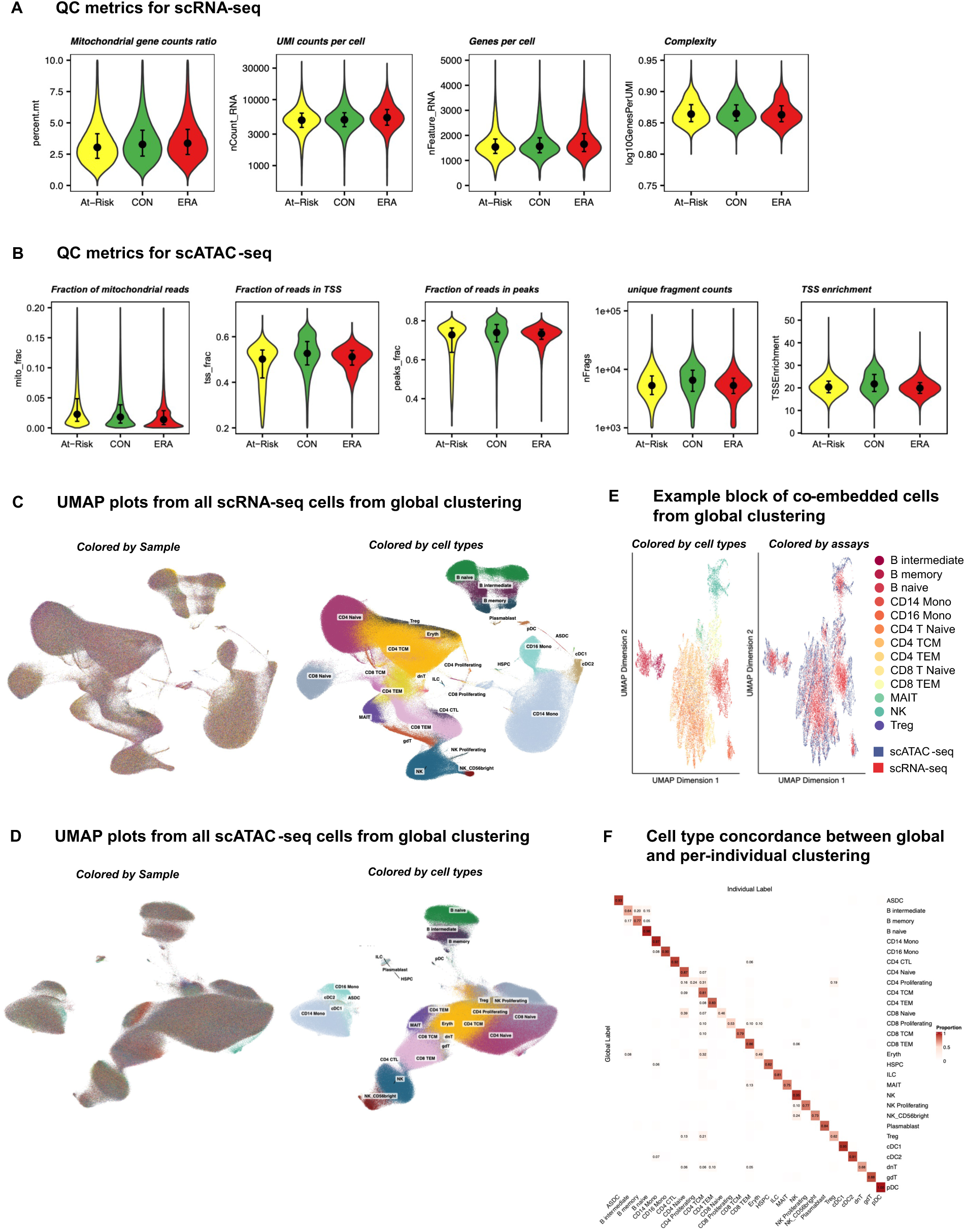
Quality control (QC) and global integration benchmarking. **(A)** Violin plots of QC metrics for scRNA-seq across At-Risk, CON, and ERA cohorts, including percent of mitochondrial gene reads, number of transcripts per cell, number of genes per cell, and library complexity (number of genes detected per UMI). **(B)** Violin plots of QC metrics for scATAC-seq, including percent of mitochondrial gene reads, fraction of reads in TSS, fraction of reads in peaks, number of unique fragments per cell, and TSS enrichment. Median (points) and 25th and 75th quantiles (whiskers and narrow bars) are overlaid on all the violin plots. Median values are also in **Supplementary Table S2**. **(C, D)** UMAP projections of scRNA-seq (**C**) and scATAC-seq (**D**) Cells generated via a standard pooled global clustering, colored by samples (left) and cell types (right). scRNA-seq cells are diffused evenly across the sample space, demonstrating a good integration across samples without batch effect. **(E)** Example block of co-embedded cells from the global clustering pipeline, colored by cell types (left) and assay modality (right). **(F)** Annotation concordance heatmap validating the high alignment between our tailored per-individual reference mapping approach and the standard global integration workflow across all cell types.

**Fig. S2.**
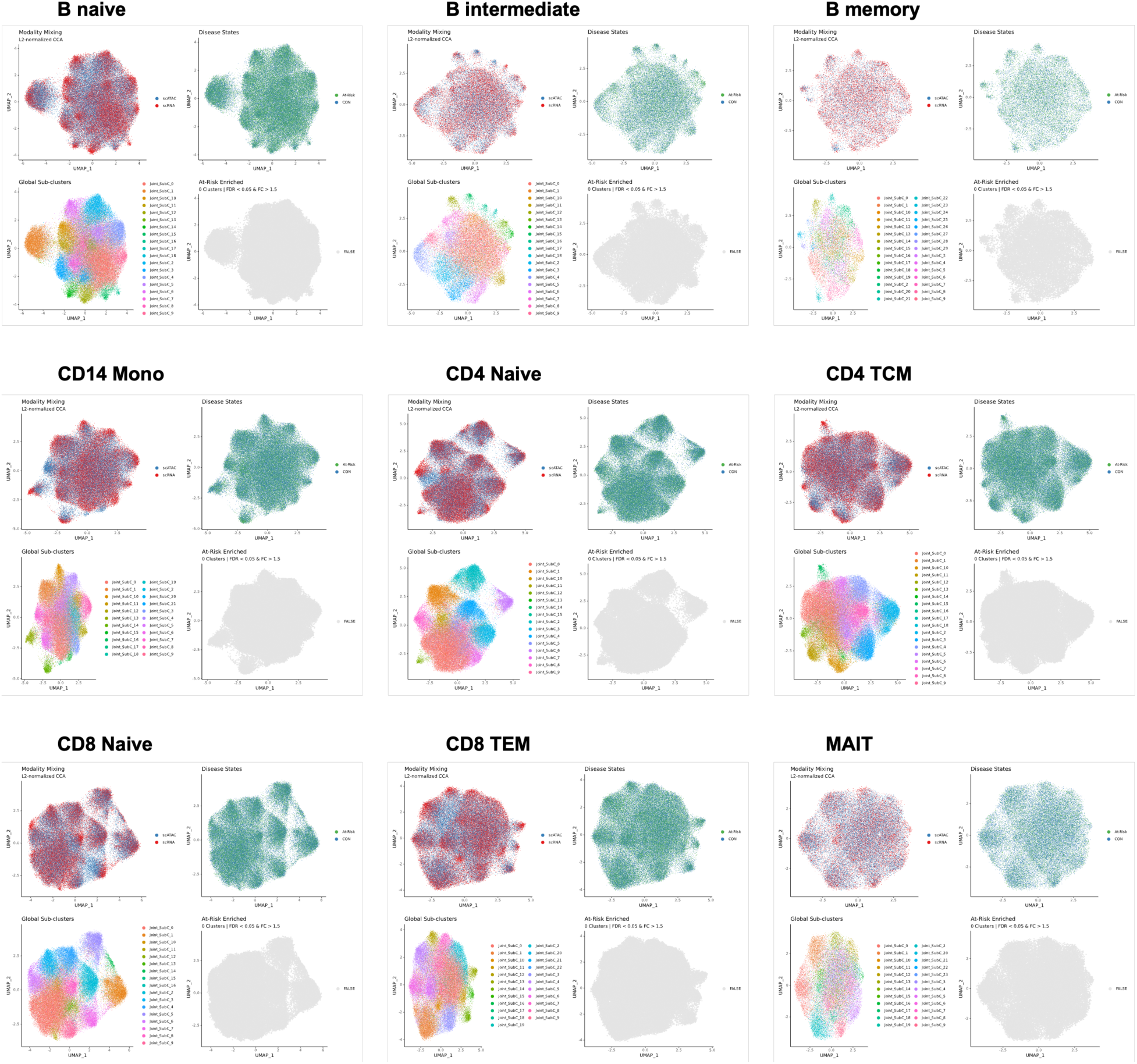
Standard global integration across all major cell lineages fails to yield At-Risk enriched sub-clusters. UMAP projections generated via a standard pooled global clustering workflow for the remaining major cell lineages (B naïve, B intermediate, B memory, CD14 Mono, CD4 T Naive, CD4 TCM, CD8 T Naive, CD8 TEM, and MAIT cells). Consistent with the NK compartment (shown in Fig. 1), the panels for each cell type depict: successful modality mixing between scATAC-seq and scRNA-seq (column 1); overlapping disease state distributions where At-Risk and CON cells are largely indistinguishable (column 2); the identification of broad global sub-clusters (column 3); and the total absence of any sub-clusters strictly enriched in the At-Risk cohort (FDR < 0.05, Fold Change > 1.5; column 4). These results demonstrate that conventional global integration broadly over-corrects patient-specific biological variance, thereby masking the multi-lineage disease signatures resolved by the per-individual analytical approach.

**Fig. S3.**
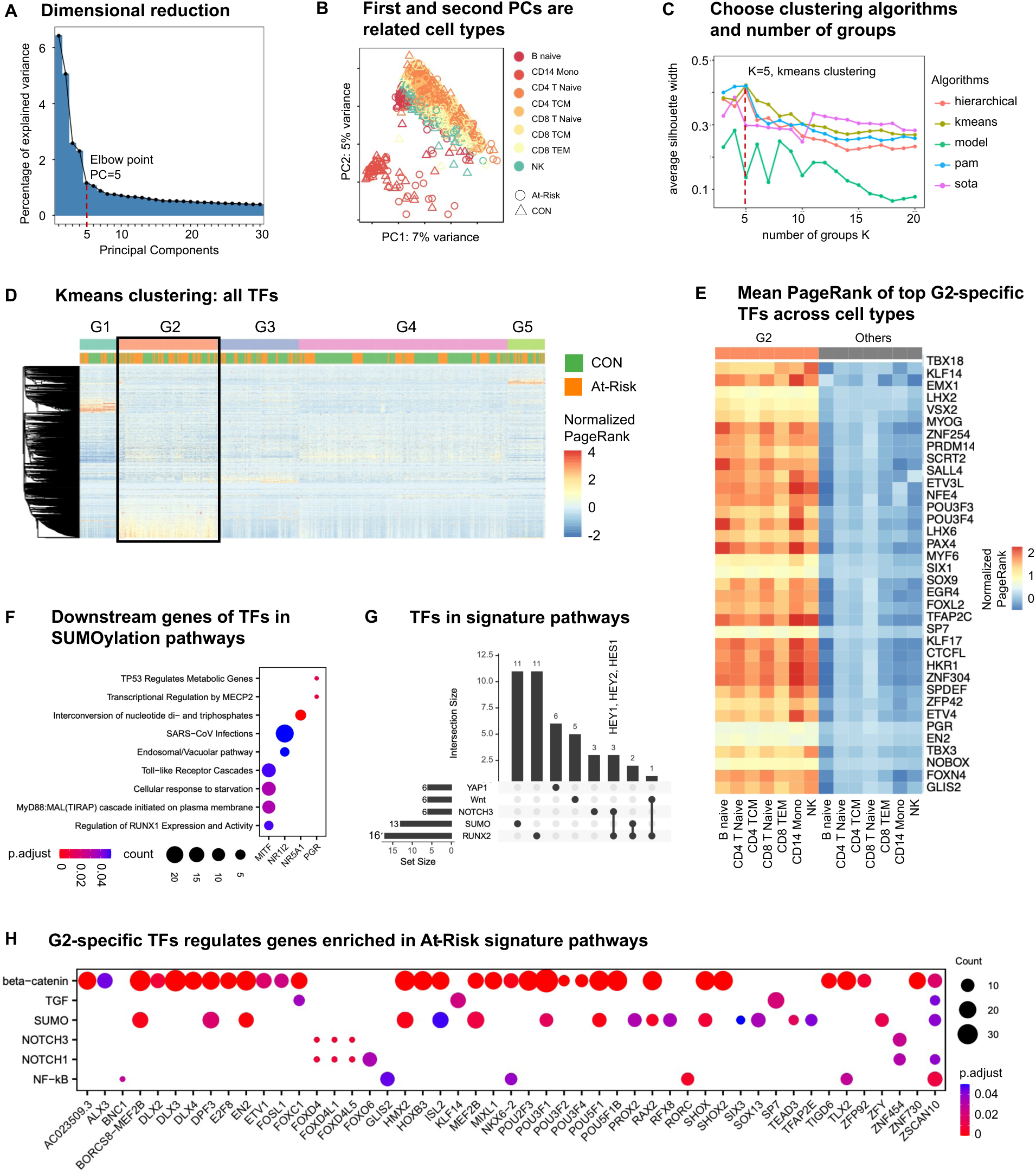
Unsupervised clustering shows distinct TF regulatory patterns. **(A)** Dimensionality reduction elbow plot of the TF PageRank score matrix, indicating PC=5 as the optimal cutoff. **(B)** First and second PCs of all clusters with color coded by cell types and shape coded by disease state. Circle represents At-Risk; triangle represents CON and square represents ERA. Each point is one cluster. Clusters are mostly separated by cell types instead of disease states. **(C)** Selecting the best clustering algorithms and number of groups K according to the silhouette metric (K=5 using K-means). **(D)** PageRank scores heatmap of all TFs across all clusters. TFs in rows (z-normalized), clusters in columns ordered by Kmeans group, and color of the cell in the matrix indicates the normalized PageRank scores with red displaying high scores. A group of TFs are significantly active in G2. **(E)** Heatmap confirming the robustness of the G2 signature by displaying the mean PageRank of the top G2-specific TFs across all major cell types, comparing G2 clusters against all other non-G2 clusters. **(F)** Dot plot of downstream target genes regulated by TFs within the enriched SUMOylation pathways. **(G)** Intersection of TFs enriched in 5 representative signature pathways. The side horizontal bars are the original size of each pathway. RUNX2 pathway shared 3 TFs with NOTCH3 pathway, 2 TFs with SUMO pathway, and 1 TF with Wnt pathway. YAP1 has its own distinct set of TFs and have no overlap with other signature pathways. **(H)** Dot plot demonstrating how G2-specific TFs regulate specific downstream genes enriched in the At-Risk signature pathways (e.g., β-catenin, TGFB, SUMO, NOTCH1/3, NF-kB).

**Fig. S4.**
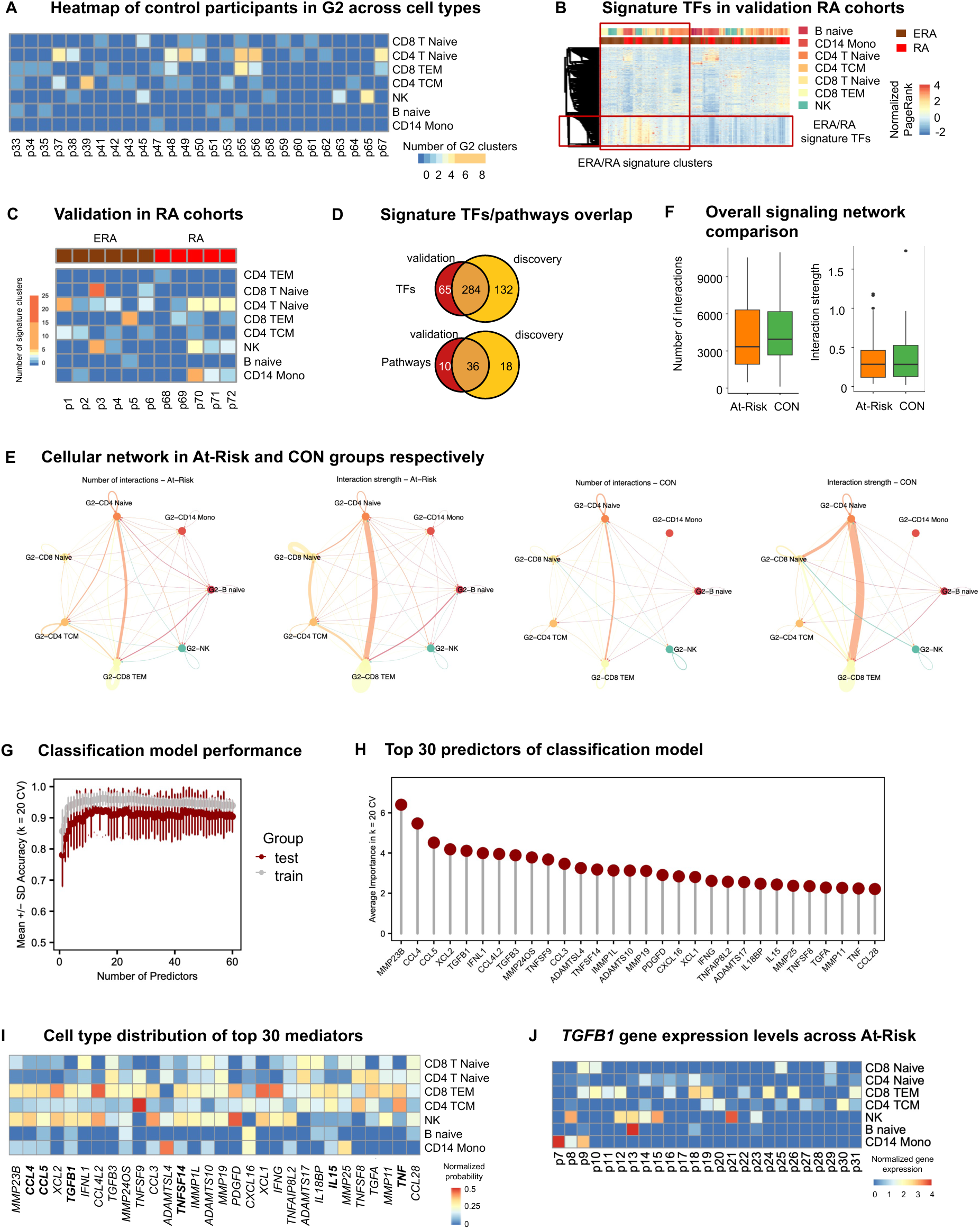
Validation of the G2 signature, communication networks, and classification model. **(A)** Heatmap showing the distribution of G2 clusters across major cell types for control (CON) participants. **(B)** Heatmap illustrating hierarchical clustering of all TFs across the early RA (ERA) and established RA validation cohorts. A group of TFs are significantly active in a group of clusters, referred to as “ERA/RA signature TFs” and “ERA/RA signature clusters”, marked by red boxes. **(C)** Heatmap depicting the highly variable distribution of signature-bearing cell types across combined validation cohorts. **(D)** Venn diagrams showing the extensive overlap of identified signature TFs (top) and enriched Reactome pathways (bottom) between the At-Risk discovery cohort and the combined RA validation cohorts. **(E)** Global cellular communication network diagrams illustrating interactions across major cell types in the At-Risk (left) and CON (right) groups. **(F)** Comparison of overall intercellular interactions between At-Risk and CON groups, which did not show any significant difference on account of number (left) and intensity (right) of interactions. **(G)** Plot showing the mean ± SD accuracy of the random forest classification model as a function of the number of gene predictors included. **(H)** Bar plot showing the average feature importance of the top 30 predictor mediators by the classification model. **(I)** Heatmap showing the diverse cell type distribution responsible for expressing the top 30 mediators. **(J)** Heatmap demonstrating the high inter-individual variability of TGFB1 gene expression across different cell lineages within the At-Risk cohort.

**Fig. S5.**
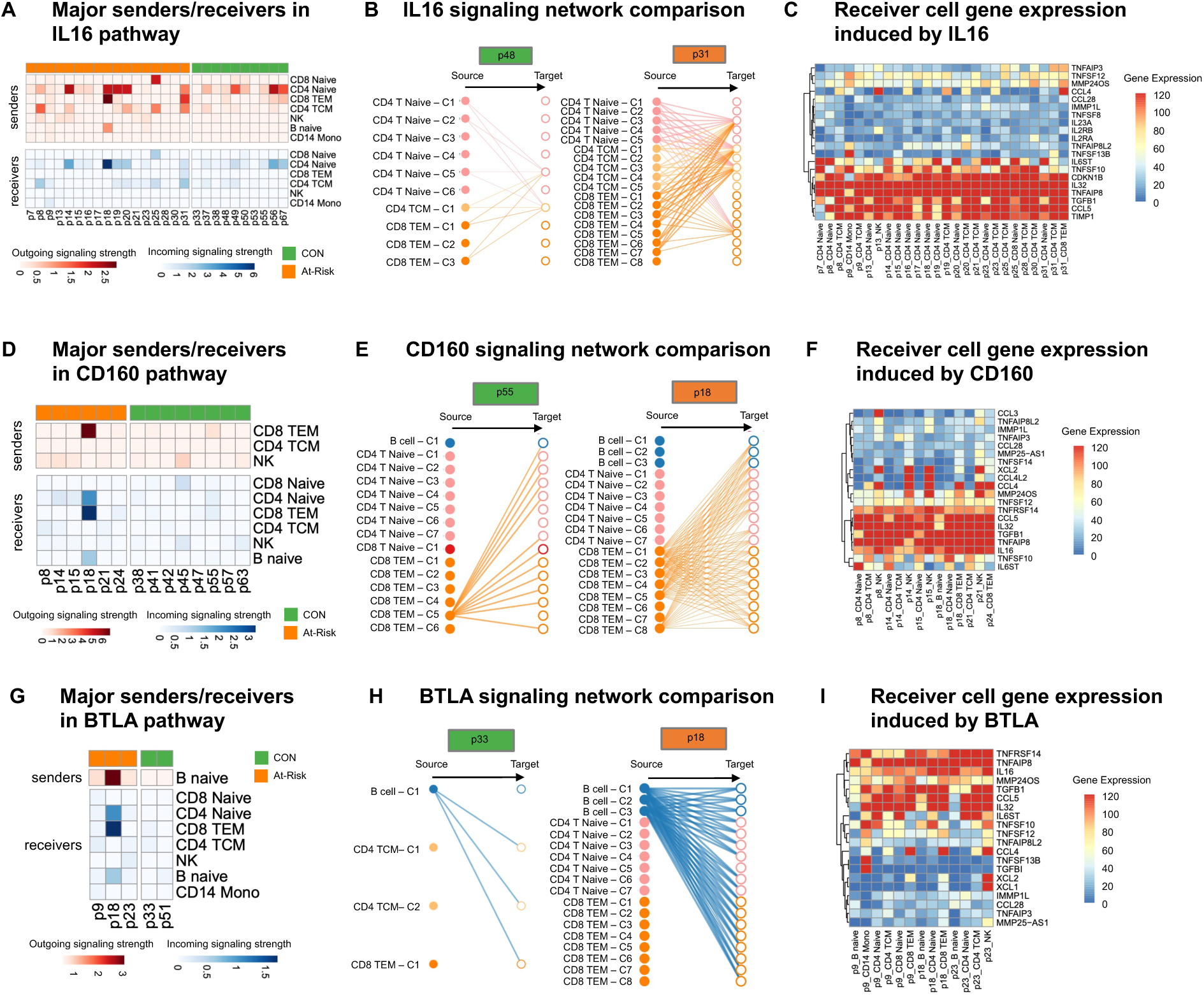
Validation of IL16, CD160, and BTLA cellular communication networks. (A, D, **G)** Representative signaling networks of IL16 **(A)**, CD160 **(D)**, and BTLA **(G)** within signature clusters in CON and At-Risk participants. At-Risk participants showed much denser and stronger interactions than controls. **(B, E, H)** Outgoing and incoming signaling strength of IL16 **(B)**, CD160 **(E)**, and BTLA **(H)** pathway across cell types in At-Risk. **(C, F, I)** Expression levels of IL16- **(C)**, CD160- **(F)**, and BTLA- **(I)** induced genes in receptor cells.

**Fig. S6.**
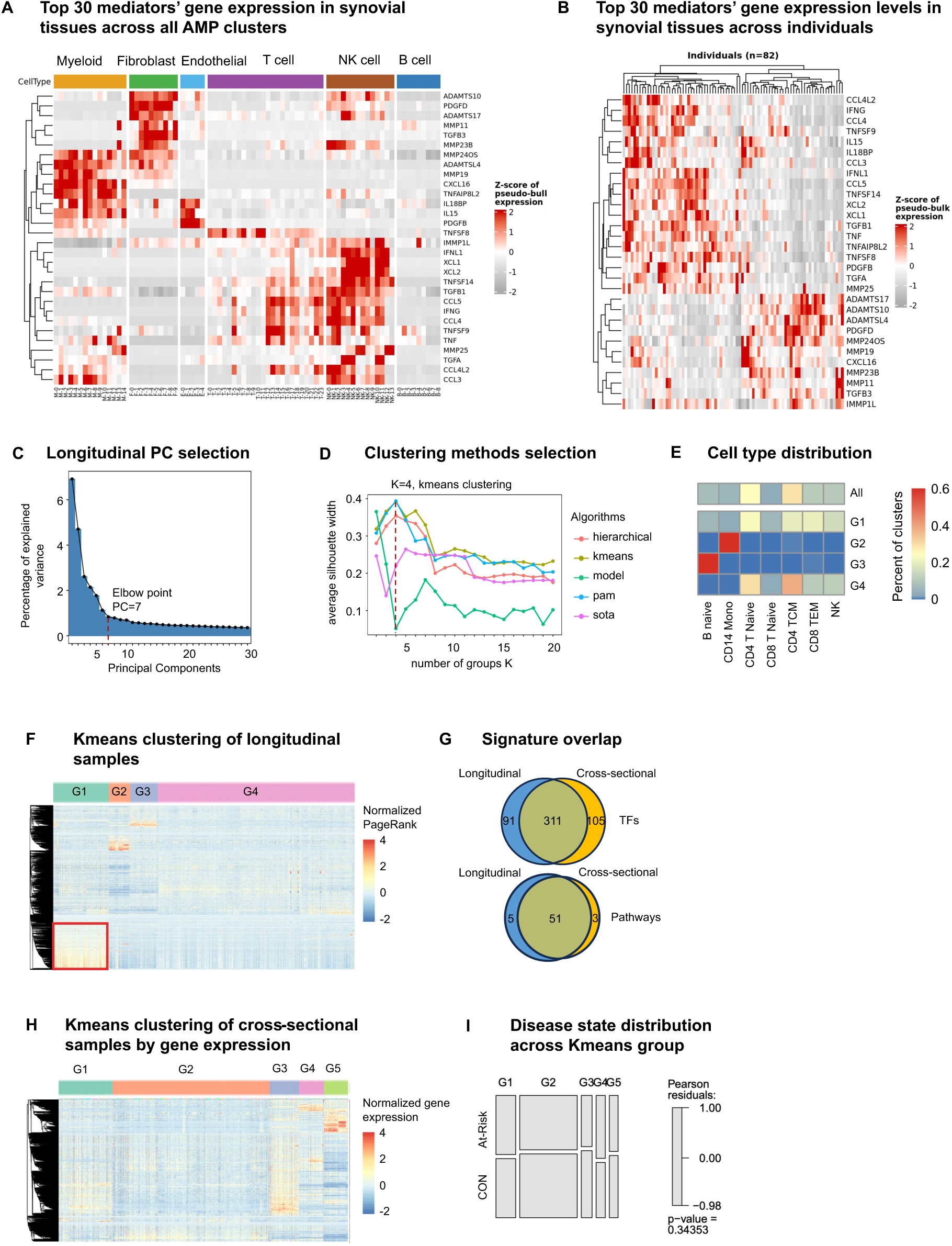
Synovial tissue validation, longitudinal clustering, and transcriptome-only benchmarking. **(A)** Heatmap depicting the expression of the top 30 PBMC-derived mediators mapped across 77 finely resolved single-cell clusters from the AMP synovial tissue dataset, explicitly highlighting tissue-resident fibroblasts alongside immune lineages. **(B)** Heatmap of the top 30 mediators’ gene expression aggregated across 82 individual RA patients in the AMP cohort, demonstrating striking patient-to-patient heterogeneity within the inflamed joint that mirrors pre-clinical peripheral blood findings. **(C)** Dimensionality reduction elbow plot for the longitudinal sample PCA, indicating PC=7. **(D)** Silhouette width analysis determining K=4 using K-means as the optimal clustering configuration for longitudinal samples. **(E)** Heatmap showing the proportional distribution of major cell types across the four longitudinal K-means groups (G1–G4). **(F)** Heatmap displaying the K-means clustering of TF PageRank scores across longitudinal samples. **(G)** Venn diagrams showing the high overlap of identified signature TFs (top) and pathways (bottom) between the cross-sectional discovery analysis and the longitudinal tracking analysis. **(H)** Heatmap demonstrating K-means clustering of cross-sectional samples utilizing just gene expression data. **(I)** Boxplot of Pearson residuals illustrating disease state distribution across the gene expression-based K-means groups from (H), demonstrating the failure of transcriptome-only data to distinctly segregate the At-Risk and CON cohorts.

## Supplementary Tables

**Supplementary Table S1.** [Summary of cohort information](https://github.com/Wang-lab-UCSD/Taiji_ALTRA/blob/main/tables/Table_S1_meta.xlsx)

**Supplementary Table S2.** [QC metrics summary for each sample](https://github.com/Wang-lab-UCSD/Taiji_ALTRA/blob/main/tables/Table_S2_qc.xlsx)

**Supplementary Table S3.** [Identified Kmeans group-specific TFs](https://github.com/Wang-lab-UCSD/Taiji_ALTRA/blob/main/tables/kmeans_specific_TFs_table.csv)

**Supplementary Table S4.** [Co-embedded cluster counts in Kmeans groups across disease states](https://github.com/Wang-lab-UCSD/Taiji_ALTRA/blob/main/tables/disease_state_freq_across_kmeans_table.csv)

**Supplementary Table S5.** [Co-embedded cluster counts in Kmeans groups across cell types](https://github.com/Wang-lab-UCSD/Taiji_ALTRA/blob/main/tables/cell_states_freq_across_kmeans_table.csv)

**Supplementary Table S6.** [Identified signature pathways with signature TFs and its representative downstream genes](https://github.com/Wang-lab-UCSD/Taiji_ALTRA/blob/main/tables/signature_TFs_and_pathways.csv)

**Supplementary Table S7.** [Curated pathogenic gene list](https://github.com/Wang-lab-UCSD/Taiji_ALTRA/blob/main/tables/gene_list_v2_rna_no_receptors.csv)

**Supplementary Table S8.** [Pathway enrichment statistics for the identified Kmeans group-specific TFs (p.adjust<0.05)](https://github.com/Wang-lab-UCSD/Taiji_ALTRA/blob/main/tables/pathway_table.csv)

## Supplementary Methods

### Genomic data acquisition

#### Sample preparation

Blood was drawn into BD NaHeparin vacutainer tubes (for PBMC; BD #367874) or K2-EDTA vacutainer tubes (for plasma; BD #367863). PBMC isolation and plasma processing were started within 2 hours post draw. For PBMC isolation, the samples in NaHeparin tubes for each donor were pooled into one common pool and combined with an equivalent volume of room temperature PBS (ThermoFisher #14190235). PBMCs were isolated using Leucosep tubes (Greiner Bio-One #227290) with 15 ml of Ficoll Premium (GE Healthcare #17-5442-03). After centrifugation, the PBMCs were recovered and resuspended with 15 ml cold PBS+0.2% BSA (Sigma #A9576; “PBS+BSA”). The cells were pelleted, resuspended in 1 ml cold PBS+BSA per 15 ml whole blood processed and counted with a Cellometer Spectrum (Nexcelom) using Acridine Orange/Propidium Iodide solution. PBMCs were cryopreserved in 90% FBS (ThermoFisher #10438026) / 10% DMSO (Fisher Scientific #D12345) at a target of 5 x 10^6^ cells/ml by slow freezing in a Coolcell LX (VWR #75779-720) overnight in a -80°C freezer followed by transfer to liquid nitrogen.

For genomics assays PBMCs were removed from liquid nitrogen storage and immediately thawed in a 37°C water bath. Cells were diluted dropwise into 40 mL AIM V media (Thermo Fisher Scientific #12055091) pre-warmed to 37°C. Cells were pelleted at 400 x g, resuspended in 5 mL cold AIM V media, and recounted using a Cellometer Spectrum. 30 mL cold AIM V media was added to the cells, which were re-pelleted and resuspended to appropriate concentration for the assays.

#### scRNA-seq

scRNA-seq was performed on PBMCs as previously described^31^ *(P. C. Genge, STAR Protoc 2, 100900 (2021))*. In brief, scRNA-seq libraries were generated using a modified 10x genomics chromium 3′ single cell gene expression assay with Cell Hashing. Sample libraries were constructed across different batches, with the addition of a common control donor leukopak sample in each library as batch control. Libraries were sequenced on the Illumina Novaseq platform. Hashed 10x Genomics scRNA-seq data processing was carried out using BarWare^35^ to generate sample-specific output files.

#### scATAC-seq

##### FACS neutrophil depletion

To remove dead cells, debris, and neutrophils prior to scATAC-seq, PBMC samples were sorted by fluorescence-activated cell sorting (FACS) following established protocols^31^. Cells were incubated with Fixable Viability Stain 510 (BD, 564406) for 15 minutes at room temperature and washed with AIM V medium (Gibco, 12055091) before incubating with TruStain FcX (BioLegend, 422302) for 5 minutes on ice, followed by staining with mouse anti-human CD45 FITC (BioLegend, 304038) and mouse anti-human CD15 PE (BD, 562371) antibodies for 20 minutes on ice. After washing, cells were then sorted on a BD FACSAria Fusion with a standard viable CD45+ cell gating scheme. Neutrophils were then excluded in the final sort gate. An aliquot of each post-sort population was used to collect 50,000 events to assess post-sort purity.

##### Sample processing

Permeabilized-cell scATAC-seq was performed as described previously^31^..A 5% w/v digitonin stock was prepared stored at −20°C. To permeabilize, 1×10^6^ cells were centrifuged and resuspended in cold isotonic Permeabilization Buffer. Then they were diluted with 1 mL of isotonic Wash Buffer and centrifuged, and the supernatant was slowly removed. Cells were resuspended in chilled TD1 buffer (Illumina, 15027866) to a target concentration of 2,300-10,000 cells per μL. Cells were filtered through 35 μm Falcon Cell Strainers (Corning, 352235) before counting on a Cellometer Spectrum Cell Counter (Nexcelom) using ViaStain acridine orange/propidium iodide solution (Nexcelom, C52-0106-5).

##### Sequencing library preparation

scATAC-seq libraries were prepared following established protocol^31^. In brief, 15,000 cells were combined with TD1 buffer (Illumina, 15027866) and Illumina TDE1 Tn5 transposase (Illumina, 15027916) and incubated at 37°C for 60 minutes. A Chromium NextGEM Chip H (10x Genomics, 2000180) was loaded and a master mix was then added to each sample well. Chromium Single Cell ATAC Gel Beads v1.1 (10x Genomics, 2000210) were loaded into the chip, along with Partitioning Oil. The chip was loaded into a Chromium Single Cell Controller instrument (10x Genomics, 120270) for GEM generation. After the run, GEMs were collected and linear amplification was performed on a C1000 Touch thermal cycler.

GEMs were separated into a biphasic mixture with Recovery Agent (10x Genomics, 220016), and the aqueous phase was retained and removed of barcoding reagents using Dynabead MyOne SILANE and SPRIselect reagent bead clean-ups. Sequencing libraries were constructed as described in the 10x scATAC User Guide. Amplification was performed in a C1000 Touch thermal cycler. Final libraries were prepared using a dual-sided SPRIselect size-selection cleanup.

##### Quantification and sequencing

Final libraries were quantified using a Quant-iT PicoGreen dsDNA Assay Kit (Thermo Fisher Scientific, P7589) on a SpectraMax iD3 (Molecular Devices). Library quality and average fragment size were assessed using a Bioanalyzer (Agilent, G2939A) High Sensitivity DNA chip (Agilent, 5067-4626). Libraries were sequenced on the Illumina NovaSeq platform with the following read lengths: 51nt read 1, 8nt i7 index, 16nt i5 index, 51nt read 2.

#### Plasma proteomics

Plasma samples were run on the Olink Explore 1536 platform. Analytes from the inflammation, oncology, cardiometabolic, and neurology panels were measured. Samples were randomized across plates to achieve a balanced distribution of age and sex. Resulting data were first normalized to an extension control that was included in each sample well. Plates were then standardized by normalizing to inter-plate controls run in triplicate on each plate. Data were then intensity normalized across all samples. Final normalized relative protein quantities were reported as log2 normalized protein expression (NPX) values by Olink. Three protein analytes were repeated across each of the four panels and treated as distinct measurements: TNF, IL-6, and CXCL8. Data, including QC flags, were reviewed for overall quality prior to analysis. Samples were measured across multiple batches.

To facilitate comparisons between batches, plasma from 12 donors was obtained commercially (BioIVT; Bloodworks Northwest) and randomly interspersed among the above study samples. Samples measured in later batches were bridge normalized to the earliest batch. Bridge offsets were determined for each batch and each analyte separately by taking the median of the per-sample NPX differences between the later batch result and the earliest (reference) batch result for the 12 commercial samples. Offsets were then subtracted from the analyte measurements of all samples in the later batch to obtain the normalized NPX values.

### Taiji pipeline overview

To characterize TF activity in each pseudo-bulk cluster, we performed an integrated multi-omics analysis using the Taiji pipeline^36,37^. Taiji integrates gene expression and epigenetic modification data to build gene regulatory networks. The algorithm first predicts putative TF binding sites in each open chromatin region that mark active promoters and enhancers using motifs documented in the CIS-BP database^38^. These TFs are then linked to their target genes predicted by EpiTensor^39^. The regulatory interactions are assembled into a genetic network. Finally, the personalized PageRank algorithm is used to assess the global influences of the TFs. In the network, the node weights are determined by the z scores of gene expression levels, allocating higher ranks to the TFs that regulate more differentially expressed genes. Each edge weight is set to be proportional to the TF’s expression level, its binding site’s open chromatin peak intensity, and the motif binding affinity, thus representing the regulatory strength. Using this method, Taiji has more power than other methods that identify key regulators in individual transcriptome and chromatin accessibility and has been confirmed using simulated data, literature evidence and experimental validation in numerous studies of various biological problems^36,37,40,41^. For this dataset, the median number of nodes and edges of the networks were 17,046 and 3,002,662, respectively, including 1047 (6.14%) TF nodes. On average, each TF regulates 3417 genes, and each gene is regulated by 184 TFs.

### TF regulatory networks weighting scheme

As described in the original Taiji paper^36^, a personalized PageRank algorithm was applied to calculate the ranking scores for TFs. We first initialized the edge weights and node weights in the network. The node weight was calculated as 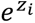, where *z_i_* is the gene’s relative expression level in cell type *i*, which is computed by applying the *z* score transformation to its absolute expression levels. The edge weight was determined by 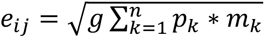., where *p* is the peak intensity, calculated as 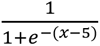, where x is −*log*_14_(*p*), represented by the p-value of the ATAC-seq peak at the predicted TF binding site, rescaled to [0, 1] by a sigmoid function; *m* is the motif binding affinity, represented by the p-value of the motif binding score, rescaled to [0, 1] by a sigmoid function; *g* is the TF expression value; *n* is the number of binding sites linked to gene *j*. Let s be the vector containing node weights and W be the edge weight matrix. The personalized PageRank score vector v was calculated by solving a system of linear equations *v* = (1 − *d*)*s* + *dWv*, where d is the damping factor (default to 0.85). The above equation can be solved in an iterative fashion, i.e., setting *v*_521_ = (1 − *d*)*s* + *dWv*_5_.

If the TFs in the same protein family share the same motifs, their PageRank scores are distinguished by their own expression levels because their motifs and the target genes are the same. If a motif is weak, the PageRank score of the TF is decided by whether these motifs occur in the open chromatin regions (measured by the peak intensity of the ATAC-seq data), the TF expression and its target expression levels. The relative difference between the PageRank scores of TFs also helps to uncover important TFs with weak motifs.

## Supplementary Notes

### Validation of Cell-Type Concordance and Data Quality

Extensive quality control was performed for both scRNA-seq and scATAC-seq modalities (**Supplementary Fig. S1A, B**). To preserve patient-specific biological variance, we utilized a per-individual, reference-based mapping approach rather than standard joint clustering. To validate this choice, we benchmarked our approach against a standard pooled global integration workflow (**Supplementary Fig. S1C-E**). As shown in the Annotation Concordance Heatmap (**Supplementary Fig. S1F**), our per-individual method yields cell populations that are highly concordant with the global consensus lineages. This demonstrates that our strategy successfully balances the preservation of patient-specific regulatory fidelity—avoiding the over-correction common in global batch integration—with robust, standardizable cell-type identification.

### Optimization of unsupervised clustering parameters

To define the optimal clustering of TF activity (PageRank scores), we systematically benchmarked dimensionality reduction and clustering algorithms. Principal Component Analysis (PCA) identified an elbow point at PC=5 (**Supplementary Fig. S3A**), which our variance analysis confirmed primarily captured robust cell-type identity (**Supplementary Fig. S3B**). We then performed Silhouette analysis to evaluate multiple algorithms (K-means, hierarchical, PAM, etc.) across various number of groups (K). K-means consistently yielded the highest performance scores, with K=5 providing the optimal average silhouette width (**Supplementary Fig. S3C**).

### Advantages of Taiji framework and Multi-Omics Integration

We measured single cell chromatin accessibility and transcriptomic profiles of PBMCs of At-Risk individuals, ERA patients and controls. Rather than relying on individual technologies and datasets to understand important pathways, we used a novel integrative analysis (Taiji)^36^ to identify potential pathogenic pathways and cell types. This method uses chromatin accessibility for TF motifs and transcription of the putatively regulated genes to prioritize TFs based on their functional relevance in a particular cell.

Taiji was previously used by our group to identify critical TFs in primary fibroblast-like synoviocytes isolated from RA synovium^42^. We were able to stratify RA patients into two groups based on divergent functions of multiple TFs and pathways. Extensive biologic validation confirmed the Taiji computational predictions related to TFs like RARA and showed that individual TFs could have diametrically opposed functions in individual patients.

Using this integrative analysis, we now report that distinctive TF profiles are significantly enriched in At-Risk individuals compared to controls in multiple cell types, especially in CD8 and CD4 T cells. This is the first time that Taiji framework has been applied to integrative single cell analysis. It’s worth noting that unsupervised clustering using solely gene expression profiles failed to reveal significant cohort enrichment and demonstrates the importance of integrating multiple omics data types (**Supplementary Fig. S6H, I**). These results support the importance of PageRank over individual gene expression or open chromatin analyses to delineate TF activities and differences across cohorts.

### G2 At-Risk TF signature is enriched for pathways implicated in RA pathogenesis

A complete review of the biology of each pathway enriched in the TF signature is beyond the scope, but it is useful to point out some of the key features and genes and their potential relationship to RA.

#### Sumoylation pathway

Sumoylation plays important regulatory roles in synovial fibroblast biology including cell survival, inflammatory responses, and matrix metabolism^14^. Several SUMO related pathways including SUMOylation, SUMO E3 ligases SUMOylate target proteins, and SUMOylation of intracellular receptors were enriched in G2 (**Supplementary Table S8**). For example, several NR family members (NR1I2, NR5A1, PGR) and MITF were found in G2 (**Supplementary Table S8**). These TFs also regulate genes significantly enriched in RA-related pathways including RUNX1, Toll-like receptors, MECP2, and TP53 pathways (**Supplementary Fig. S3I**). NR1I2 regulates HLA-G, which plays an important role in RA susceptibility and regulation.

#### RUNX2 and NOTCH3 pathways

While most signature pathways possessed distinctive sets of active TFs, the RUNX2 pathway shared 37.5% TFs with other signature pathways, particularly with the NOTCH3 pathway (**Supplementary Fig. S3J** ), which suggests the interdependence between RUNX2 and other signature pathways. For instance, three TFs (HEY1, HEY2, and HES1) were identified in both NOTCH3 and RUNX2 pathways and regulate osteoblast function^43^. NOTCH genes and signaling also play a critical role in the differentiation of synovial fibroblasts into pathogenic cells^44^.

#### YAP1 pathway

Recent evidence suggests a critical regulatory role of the Hippo pathway in the RA pathogenesis. As one of the key components, YAP promotes the localization of SMAD3 in RA fibroblast-like synoviocytes and enhances aggressiveness^45^. Unlike RUNX2 and NOTCH3 pathways, YAP1 pathway did not share any TFs with other signature pathways (**Supplementary Fig. S3G**). Multiple TEAD family members were also identified as important in the YAP1/TAZ pathway by binding and promoting gene expression. Previous studies indicate that inhibiting YAP-TEAD interaction reduces RA synoviocytes invasion^16^.

#### β-catenin pathway

Although only a few TFs were enriched in β-catenin pathways, many of the TFs including DLX3, MEF2B, POU3F1, RAX2, TLX2 from different families had regulatees involved in β-catenin-related pathways (**Supplementary Fig. S3H**). For example, PAX4 regulates many functional genes including BCL9L, CARD11, AURKB, NR3C1, and BIRC2 that are significantly enriched in the signature pathways.

Other pathways. Besides of the signature pathways which are well known to be involved in RA pathogenesis, we also found some novel pathways of potential interest in RA (**Supplementary Table S8**). For example, regulation of beta-cell development was ranked as one of the top G2-specific pathways, in which many TFs from HNF and NKX families are involved. Other examples are transcriptional regulation of pluripotent stem cells pathway and regulation of gene expression in late-stage pancreatic bud precursor cells pathway.

G4 is significantly enriched in the CON cohort (32% higher in CON, p-value < 0.0001; Chi-squared test) (**Fig. 2D**). β-catenin, Wnt, and RUNX related pathways were enriched in both G2 and G4 (**Fig. 2E**), suggesting potential multifaceted roles for specific signature pathways across different participant groups. However, the genes implicated in G4 were distinct from G2. For example, the former was enriched in the formation of the β-catenin:TCF transactivating complex while latter was associated with the deactivation of the β-catenin transactivating complex. This is consistent with the observations that Wnt/β-catenin signaling can exert either anti-inflammatory and proinflammatory functions depending on the context^46^. Similarly, RUNX3, which is also a pathway associated with CON, is chondroprotective in preclinical models of arthritis^47^.

### Consistency of G2 signature across cell types

To confirm the pan-lineage nature of the G2 signature, we evaluated the mean PageRank of the top G2-specific TFs across all major cell types. As shown in **Supplementary Fig. S3E**, these top signature TFs are consistently highly active within the G2 group across diverse cellular compartments compared to non-G2 clusters, confirming that this regulatory state transcends individual cell lineages.

### Combined analysis of validation RA cohorts

We confirmed the persistence of the At-Risk signature in patients by combining two clinical RA cohorts. Hierarchical clustering of TF PageRank scores in the validation cohorts (**Supplementary Fig. S4B**) revealed a consistent regulatory signature identical to the pre-clinical state. Importantly, the distribution of signature-bearing cell types remained highly heterogeneous across individual RA patients (**Supplementary Fig. S4C**), mirroring the cellular plasticity seen in the At-Risk cohort. Furthermore, Venn diagrams demonstrate a massive and highly significant overlap in both the identified signature TFs and their enriched Reactome pathways between the At-Risk and established RA cohorts (**Supplementary Fig. S4D**), confirming the temporal stability of this core pathogenic module.

### Classification model performance and mediator source heterogeneity

In developing the random forest classification model, we evaluated prediction accuracy as a function of the number of gene predictors. Model accuracy monotonically increased with more predictors and reached a stable plateau at approximately 30 predictors (**Supplementary Fig. S4G**). While this core set of 30 pathogenic mediators was robustly shared across the At-Risk cohort, the cellular sources producing them were highly variable. For instance, evaluating the expression of TGFB1 across all At-Risk individuals (**Supplementary Fig. S4J**) clearly illustrates that the predominant cellular source of this single key mediator shifts dramatically from patient to patient (e.g., driven by CD8 TEM in some patients, and B naive or Monocytes in others).

### Synovial Tissue Mapping (AMP Dataset)

To directly address tissue-level relevance, we comprehensively mapped our top 30 blood-derived predictors against 77 finely resolved synovial sub-clusters from the AMP Phase 2 dataset (**Supplementary Fig. S6A**). Importantly, this analysis explicitly incorporated tissue-resident stromal populations. We discovered highly compartmentalized expression: while inflammatory chemokines (e.g., CCL5, IFNG) were predominantly expressed by tissue-infiltrating T and NK cells, a distinct module of our blood-derived predictors—including tissue remodeling enzymes (MMPs, ADAMTS family) and specific growth factors (PDGFD, TGFB3)—was intensely and specifically expressed by synovial fibroblasts. This strongly reinforces that the systemic signature seeds coordinated, multi-cellular crosstalk with stromal cells in the target joint (**Supplementary Fig. S6B**).

### Longitudinal Clustering Selection

For the longitudinal samples, we repeated our optimization pipeline. PCA selection identified an elbow at PC=7 (**Supplementary Fig. S6C**), and Silhouette analysis determined K=4 as the optimal cluster number using K-means (**Supplementary Fig. S6D**). The resulting multi-lineage clusters (**Supplementary Fig. S6E, F**) revealed dynamic TF activity states that significantly overlapped (60% of TFs, 86% of pathways) with the pathogenic signatures identified in the cross-sectional discovery dataset (**Supplementary Fig. S6G**).

## References

1. Gravallese, E. M. & Firestein, G. S. Rheumatoid Arthritis - Common Origins, Divergent Mechanisms. N. Engl. J. Med. 388, (2023).

2. Holers, V. M. et al. Mechanism-driven strategies for prevention of rheumatoid arthritis. Rheumatology & autoimmunity 2, 109–119 (2022).

3. Holers, V. M. et al. Rheumatoid arthritis and the mucosal origins hypothesis: protection turns to destruction. Nat. Rev. Rheumatol. 14, 542–557 (2018).

4. Deane, K. D. et al. A Phase 2 Trial of Hydroxychloroquine in Individuals at Risk for Rheumatoid Arthritis. Arthritis Rheumatol 78, 809–820 (2026).

5. Rech, J. et al. Abatacept inhibits inflammation and onset of rheumatoid arthritis in individuals at high risk (ARIAA): a randomised, international, multicentre, double-blind, placebo-controlled trial. Lancet 403, 850–859 (2024).

6. van Boheemen, L. et al. Atorvastatin is unlikely to prevent rheumatoid arthritis in high risk individuals: results from the prematurely stopped STAtins to Prevent Rheumatoid Arthritis (STAPRA) trial. RMD open 7, e001591 (2021).

7. Gerlag, D. M. et al. Effects of B-cell directed therapy on the preclinical stage of rheumatoid arthritis: the PRAIRI study. Ann. Rheum. Dis. 78, 179–185 (2019).

8. Krijbolder, D. I. et al. Intervention with methotrexate in patients with arthralgia at risk of rheumatoid arthritis to reduce the development of persistent arthritis and its disease burden (TREAT EARLIER): a randomised, double-blind, placebo-controlled, proof-of-concept trial. Lancet 400, 283–294 (2022).

9. Weinand, K. et al. The chromatin landscape of pathogenic transcriptional cell states in rheumatoid arthritis. Nature Communications 15, 4650 (2024).

10. Zhang, F. et al. Defining inflammatory cell states in rheumatoid arthritis joint synovial tissues by integrating single-cell transcriptomics and mass cytometry. Nat Immunol 20, 928–942 (2019).

11. Zhang, F. et al. Deconstruction of rheumatoid arthritis synovium defines inflammatory subtypes. Nature 623, 616–624 (2023).

12. He, Z. et al. Progression to rheumatoid arthritis in at-risk individuals is defined by systemic inflammation and by T and B cell dysregulation. Sci Transl Med 17, eadt7214 (2025).

13. Intlekofer, A. M. et al. Effector and memory CD8+ T cell fate coupled by T-bet and eomesodermin. Nat. Immunol. 6, 1236–1244 (2005).

14. Dehnavi, S. et al. The role of protein SUMOylation in rheumatoid arthritis. J. Autoimmun. 102, 1–7 (2019).

15. Di Chen, Dongyeon J Kim, Jie Shen, Zhen Zou, Regis J O’Keefe. Runx2 plays a central role in Osteoarthritis development. Journal of Orthopaedic Translation 23, 132–139 (2020).

16. Caire, R. et al. YAP/TAZ: Key Players for Rheumatoid Arthritis Severity by Driving Fibroblast Like Synoviocytes Phenotype and Fibro-Inflammatory Response. Front. Immunol. 12, 791907 (2021).

17. Zhuang, Y. et al. A narrative review of the role of the Notch signaling pathway in rheumatoid arthritis. Annals of Translational Medicine 10, 371–371 (2022).

18. Chen, S. et al. Wnt/β-catenin signaling pathway promotes abnormal activation of fibroblast-like synoviocytes and angiogenesis in rheumatoid arthritis and the intervention of Er Miao San. Phytomedicine 120, 155064 (2023).

19. Galea, C. A., Nguyen, H. M., George Chandy, K., Smith, B. J. & Norton, R. S. Domain structure and function of matrix metalloprotease 23 (MMP23): role in potassium channel trafficking. Cell. Mol. Life Sci. 71, 1191–1210 (2013).

20. Serum proteomic analysis identifies interleukin 16 as a biomarker for clinical response during early treatment of rheumatoid arthritis. Cytokine 78, 87–93 (2016).

21. Iwasaki, T. et al. Monocyte-derived transcriptomes explain the ineffectiveness of abatacept in rheumatoid arthritis. Arthritis Res Ther 26, 1 (2024).

22. Choi, E. et al. Joint-specific rheumatoid arthritis fibroblast-like synoviocyte regulation identified by integration of chromatin access and transcriptional activity. JCI Insight 9, e179392 (2024).

23. Inamo, J. et al. Deep immunophenotyping reveals circulating activated lymphocytes in individuals at risk for rheumatoid arthritis. J Clin Invest 135, e185217 (2025).

24. Moreland, L. W. et al. Double-blind, placebo-controlled multicenter trial using chimeric monoclonal anti-CD4 antibody, cM-T412, in rheumatoid arthritis patients receiving concomitant methotrexate. Arthritis Rheum 38, 1581–1588 (1995).

25. He, S. et al. A longitudinal cohort study uncovers plasma protein biomarkers predating clinical onset and treatment response of rheumatoid arthritis. Nat Commun 16, 6692 (2025).

26. Prideaux, E. B. et al. Epigenetic trajectory predicts development of clinical rheumatoid arthritis in ACPA+ individuals: Targeting Immune Responses for Prevention of Rheumatoid Arthritis (TIP-RA). bioRxiv 2024.10.15.618490 (2025) doi:10.1101/2024.10.15.618490.

27. Joehanes, R. et al. Epigenetic Signatures of Cigarette Smoking. Circ. Cardiovasc. Genet. 9, 436–447 (2016).

28. James, E. A. et al. Multifaceted immune dysregulation characterizes individuals at-risk for rheumatoid arthritis. Nat. Commun. 14, 7637 (2023).

29. Warren, R. B. et al. Long-Term Efficacy and Safety of Bimekizumab and Other Biologics in Moderate to Severe Plaque Psoriasis: Updated Systematic Literature Review and Network Meta-analysis. Dermatol Ther (Heidelb) 14, 3133–3147 (2024).

30. Aletaha, D. et al. 2010 Rheumatoid arthritis classification criteria: an American College of Rheumatology/European League Against Rheumatism collaborative initiative. Arthritis Rheum. 62, 2569–2581 (2010).

31. Swanson, E. et al. Simultaneous trimodal single-cell measurement of transcripts, epitopes, and chromatin accessibility using TEA-seq. Elife 10, e63632 (2021).

32. Yuhan Hao, Stephanie Hao, Erica Andersen-Nissen, William M. Mauck, Shiwei Zheng, Andrew Butler, Maddie J. Lee, Aaron J. Wilk, Charlotte Darby, Michael Zager, Paul Hoffman, Marlon Stoeckius, Efthymia Papalexi, Eleni P. Mimitou, Jaison Jain, Avi Srivastava, Tim Stuart, Lamar M. Fleming, Bertrand Yeung, Angela J. Rogers, Juliana M. McElrath, Catherine A. Blish, Raphael Gottardo, Peter Smibert, Rahul Satija. Integrated analysis of multimodal single-cell data. Cell 184, 3573–3587 (2021).

33. Zhang, B. et al. CD127 imprints functional heterogeneity to diversify monocyte responses in inflammatory diseases. J Exp Med 219, e20211191 (2022).

34. Avila Cobos, F., Alquicira-Hernandez, J., Powell, J. E., Mestdagh, P. & De Preter, K. Benchmarking of cell type deconvolution pipelines for transcriptomics data. Nat Commun 11, 5650 (2020).

35. Swanson, E., Reading, J., Graybuck, L. T. & Skene, P. J. BarWare: efficient software tools for barcoded single-cell genomics. BMC Bioinformatics 23, 106 (2022).

36. Zhang, K., Wang, M., Zhao, Y. & Wang, W. Taiji: System-level identification of key transcription factors reveals transcriptional waves in mouse embryonic development. Sci Adv 5, eaav3262 (2019).

37. Yu, B. et al. Epigenetic landscapes reveal transcription factors that regulate CD8 T cell differentiation. Nature Immunology 18, 573–582 (2017).

38. Weirauch, M. T. et al. Determination and inference of eukaryotic transcription factor sequence specificity. Cell 158, (2014).

39. Zhu, Y. et al. Constructing 3D interaction maps from 1D epigenomes. Nat. Commun. 7, 10812 (2016).

40. Liu, C. et al. Systems-level identification of key transcription factors in immune cell specification. PLoS Comput. Biol. 18, e1010116 (2022).

41. Chung, H. K. et al. Atlas-guided discovery of transcription factors for T cell programming. Nature 651, 1077–1087 (2026).

42. Ainsworth, R. I. et al. Systems-biology analysis of rheumatoid arthritis fibroblast-like synoviocytes implicates cell line-specific transcription factor function. Nat. Commun. 13, 1–11 (2022).

43. Hilton, M. J. et al. Notch signaling maintains bone marrow mesenchymal progenitors by suppressing osteoblast differentiation. Nat. Med. 14, 306–314 (2008).

44. Wei, K. et al. Notch signaling drives synovial fibroblast identity and arthritis pathology. Nature 582, 259–264 (2020).

45. Bottini, A. et al. PTPN14 phosphatase and YAP promote TGFβ signalling in rheumatoid synoviocytes. Ann. Rheum. Dis. 78, 600–609 (2019).

46. Ma, B. & Hottiger, M. O. Crosstalk between Wnt/β-Catenin and NF-κB Signaling Pathway during Inflammation. Front. Immunol. 7, 221254 (2016).

47. Nagata, K. et al. Runx2 and Runx3 differentially regulate articular chondrocytes during surgically induced osteoarthritis development. Nat. Commun. 13, 6187 (2022).

